# Protease-mediated processing of Argonaute proteins controls small RNA association

**DOI:** 10.1101/2020.12.09.417253

**Authors:** Rajani Kanth Gudipati, Kathrin Braun, Foivos Gypas, Daniel Hess, Jan Schreier, Sarah H. Carl, René F. Ketting, Helge Großhans

## Abstract

Small RNA pathways defend the germlines of animals against selfish genetic elements and help to maintain genomic integrity. At the same time, their activity needs to be well-controlled to prevent silencing of ‘self’ genes. Here, we reveal a proteolytic mechanism that controls endogenous small interfering (22G) RNA activity in the *Caenorhabditis elegans* germline to protect genome integrity and maintain fertility. We find that WAGO-1 and WAGO-3 Argonaute (Ago) proteins are matured through proteolytic processing of their unusually proline-rich N-termini. In the absence of DPF-3, a P-granule-localized N-terminal dipeptidase orthologous to mammalian DPP8/9, processing fails, causing a change of identity of 22G RNAs bound to these WAGO proteins. Desilencing of repeat- and transposon-derived transcripts, DNA damage and acute sterility ensue. These phenotypes are recapitulated when WAGO-1 and WAGO-3 are rendered resistant to DFP-3-mediated processing, identifying them as critical substrates of DPF-3. We conclude that N-terminal processing of Ago proteins regulates their activity and promotes discrimination of self from non-self by ensuring association with the proper complement of small RNAs.

**Graphical Abstract: The role of DPF-3 in the fertility of the animals:** 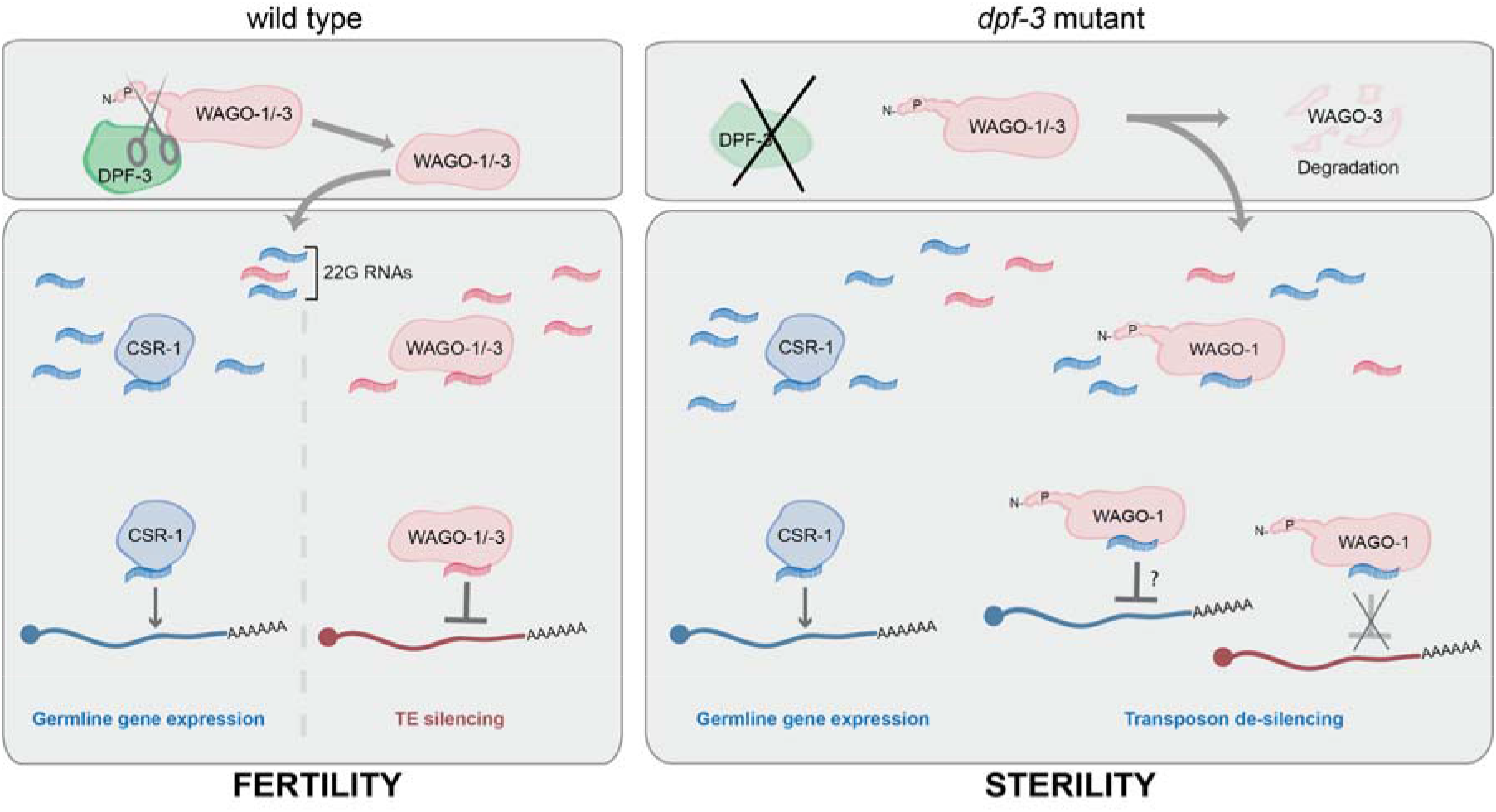

In wild type animals, the WAGO-1 and WAGO-3 Argonaute proteins are produced as immature pro-proteins with N-termini (N) that are unusually rich in prolines (P). N-terminal processing by DPF-3 is required for loading of the proper small RNA cargo and stabilization of WAGO-3. Accordingly, loss of this processing activity causes desilencing of transposable elements (TE), cell death and sterility.

## Introduction

Members of the DPPIV serine protease family have important roles in animal pathophysiology. Thus, DPPIV itself functions as a regulator of glucose homeostasis, and its pharmaceutical inhibition is clinically exploited to treat type II diabetes (Deacon, 2019), while the paralogous DPP8 and DPP9 play a role in immune system function and malignancies, inflammation, and beyond (Zhang *et al*., 2013). DPPIV family (DPF) serine proteases cleave off two amino acid-long peptides of the sequence Xaa-Pro or Xaa-Ala from the N-terminus (Mentlein, 1999; Keane *et al*., 2011). Xaa may be methionine as the initiating amino acid in translation, but this amino acid is frequently co-translationally removed in eukaryotes (Wingfield, 2017) and DPF proteases accept any amino acid in this position. The initial cleavage by DPPIV proteases initiates further degradation or renal clearance of the substrates (Mentlein, Gallwitz and Schmidt, 1993; Justa-Schuch *et al*., 2016). Several other regulatory roles have been proposed for DPPIV proteases (Wilson *et al*., 2016) but none have been verified yet. Here we uncover the function of a DPF protease in protein maturation and functional modulation.

To explore the functions of this protease family more broadly, we studied them in *C. elegans*, where seven genes encode DPF proteins of which six, DPF-1 through DPF-6, exhibit conservation of the catalytic triad (fig. S1A,B). Through targeted mutagenesis of these six genes, we uncovered a role of DPF-3, orthologous to mammalian DPP8/9 in promoting fertility by controlling the activity of small RNA-based silencing pathways.

Small RNA-mediated antisense silencing mechanisms have evolved to control the activity of selfish genetic elements such as transposable elements (TE) and repeats, which, unchecked, are a threat to genome integrity in animal germlines (Gu *et al*., 2009). In *C. elegans*, a subset of Argonaute (Ago) proteins known as WAGOs (<u>W</u>orm <u>A</u>r<u>GO</u>nautes) utilize endogenous small interfering RNAs termed 22G RNAs (because of their length and 5’-terminal nucleotide bias) to target TEs. Among several WAGOs with presumably partially redundant function (Yigit *et al*., 2006), WAGO-1, possibly along with the highly similar WAGO-3, is particularly important for 22G RNA accumulation and TE silencing (Vastenhouw *et al*., 2003; Gu *et al*., 2009; de Albuquerque, Placentino and Ketting, 2015).

The WAGO pathway, like other defense pathways targeting self-similar elements, faces a conundrum: It should have the widest possible target spectrum yet silencing of self (endogenous) genes needs to be avoided. The sequence diversity of 22G RNAs combined with their ability to recognize individual target RNAs with high specificity through base pairing makes them potentially well suited to balancing these distinct needs. However, it requires additional mechanisms to define the proper cellular complement of 22G RNAs. In part, these appear to be contributed by two other classes of small RNAs, namely 21U RNAs bound by PRG-1 (Batista *et al*., 2008; de Albuquerque, Placentino and Ketting, 2015; Phillips *et al*., 2015) and 26G RNAs bound by ALG-3/-4 and ERGO-1 Ago proteins (Vasale *et al*., 2010; Conine *et al*., 2010). PRG-1/ALG-3/-4/ERGO-1 binding turns RNAs into substrates of RNA-dependent RNA Polymerases (RdRPs), which then synthesize 22G RNAs (Almeida, de Jesus Domingues and Ketting, 2019). However, since 26G and 21U RNAs appear capable of scanning much, or even all, of the germline transcriptome themselves (Seth *et al*., 2013; Shen *et al*., 2018; Wedeles, Wu and Claycomb, 2014), further mechanisms are needed to prevent unintended activity of WAGOs on self transcripts.

Previous work identified CSR-1-dependent upregulation of endogenous germline genes as a mechanism important for insulating them from inappropriate silencing by WAGOs (Conine *et al*., 2013; Seth *et al*., 2013; Wedeles, Wu and Claycomb, 2013). Paradoxically, however, this Ago protein also relies on 22G RNAs to recognize its targets (Claycomb *et al*., 2009). Hence, rather than solving the challenge of achieving proper self vs. non-self discrimination, the opposing activities of WAGOs and CSR-1 require mechanisms that can specify the distinct pools of 22G RNAs with which they can associate.

Here, we reveal that DPF-3 processes WAGO-1 and WAGO-3 to control 22G RNA accumulation. Loss of *dpf-3* causes failure of small RNA sorting to WAGO-1 and destabilization of WAGO-3, resulting in TE desilencing, DNA damage, loss of sperm and their progenitors, and acute sterility.

## Results

### Loss of function of DPF-3 leads to a decrease in brood size

To explore the functions of DPF proteins in a whole-animal context, we created deletions in each of the uncharacterized *C. elegans dpf-1~6* genes (fig. S1C). Most deletions appeared well tolerated, with mutant animals looking overtly wild-type under standard culture conditions. However, we noticed unfertilized oocytes on the plates with *dpf-3(xe68)* null mutant animals (hereafter referred to as *dpf-3Δ*). Moreover, at 25°C, these animals had a noticeably reduced brood relative to wild-type animals (Fig. 1A).

**Fig. 1.**
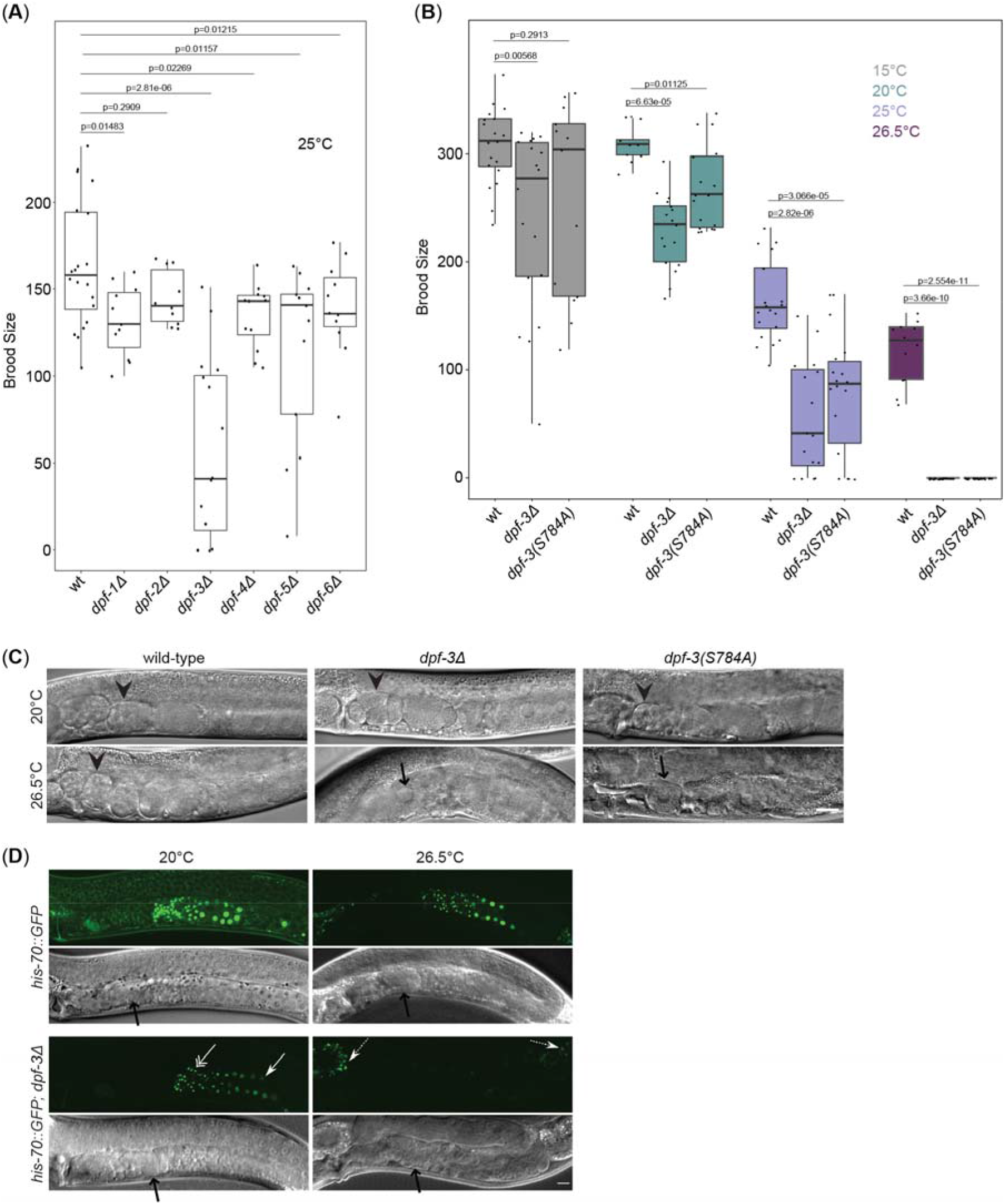
Loss of *dpf-3* function leads to fertility defects and sterility caused by defective spermatogenesis. (**A**) Box plot of the brood sizes of wild-type (wt) and indicated *dpf* mutant animals grown at 25°C and (**B**) of wt and *dpf-3* mutant animals grown at different temperatures as indicated with a color code. Each dot represents an individual worm. The same data are used for wt and *dpf-3Δ* animals at 25°C in both plots. p values are calculated using a Wilcoxon test. (**C**) DIC images of wild-type or *dpf-3* mutant animals grown at the indicated temperatures. Arrowheads indicate embryos, arrows unfertilized oocytes in the uterus. Scale bar is 10 μm. (**D**) Confocal images of L4-stage animals expressing *his-70::gfp*. A double headed white arrow indicates spermatids, single headed white solid arrows sperm progenitor cells, a dashed white arrow autofluorescence. Black arrows in the DIC images indicate spermathecae. Scale bar is 10 μm.

DPF-3 is orthologous to mammalian DPP8/9, for which catalytic activity depends on the catalytic triad, comprised of Ser, His and Asp (Ajami *et al*., 2003). To understand if DPF-3 catalytic activity is required for reproduction, we used genome editing to change the catalytic Ser 784 to Ala (S784A), generating *dpf-3(xe71[S784A])* mutant animals. In the following, we will refer to this strain, in which DPF-3 is additionally FLAG-HA epitope-tagged, as *dpf-3(S784A)*; we validated loss of catalytic activity as described in a later section.

Although DPF-3 protein levels were comparable between mutant and wild-type animals (fig. S1D), the mutation also caused a notable decrease in fertility (Fig. 1B). Moreover, for both the catalytic dead and the null mutant allele of *dpf-3*, we observed a temperature-dependence of the phenotype: The brood size of mutant animals decreased with increasing temperature and complete sterility occurred with both alleles at 26.5°C (Fig. 1B). Individual mutations of the other two residues of the catalytic triad resulting in *dpf-3(D861A)* or *dpf-3(H893A)* also caused fully penetrant sterility at 26.5°C (fig. S1E) and mutant animals could not be propagated at this temperature. Finally, introduction of a functional copy of *dpf-3* in *dpf-3Δ* background (*dpf-3* rescue) restored fertility (fig. S1E, F), confirming specificity of the observed phenotype. We conclude that DPF-3, and in particular its catalytic activity, is required for animal fertility.

### Spermatogenesis is affected in *dpf-3* mutant animals

To identify the cause of sterility, we examined wild-type and *dpf-3* mutant animals by Differential Interference Contrast (DIC) microscopy. When grown at 20°C, all animals contained fertilized eggs and multicellular embryos in the uterus. However, at 26.5°C, this was true for wild-type, but not for *dpf-3* mutant animals. Instead of embryos, *dpf-3* mutant animals carried unfertilized oocytes in the uterus (Fig. 1C).

To assess whether this apparent fertilization defect was a consequence of sperm or oocyte defects, or both, we performed crosses between hermaphrodites and males. When we mated *dpf-3Δ* mutant hermaphrodites grown at 26.5°C with wild-type males grown at 20°C, we could readily observe viable progeny (fig. S1G). Moreover, *dpf-3Δ* males grown at 20°C could sire progeny with the *dpf-3Δ* hermaphrodites grown at 26.5°C (fig. S1G), while *dpf-3Δ* males grown at 26.5° could not (n>100). Hence, *dpf-3* causes a male fertility defect at the elevated temperature.

To explore this further, we examined hermaphrodites at the L4 stage, when spermatogenesis but not oogenesis is happening to visualize sperm progenitor cells and mature sperm using a *his-70::GFP* reporter specifically expressed in these cells (Delaney *et al*., 2018). Both types of cells were readily detectable in wild-type animals irrespective of growth temperature. By contrast, in *dpf-3Δ* mutant animals, they were only visible at 20°C but undetectable at 26.5°C (Fig. 1D), revealing that DPF-3 supports sperm production at elevated temperatures.

We note that wild-type males sired fewer progeny when crossed to *dpf-3* mutant than to wild-type hermaphrodites (fig. S1G). Combined with additional evidence provided below, this indicates that DPF-3 may also promote female fertility. Because *C. elegans* is a serial hermaphrodite that produces only a limited number of sperm during the last larval stage before switching to oocyte production, which continues throughout adulthood (L’Hernault, 2009), the male fertility defect may dominate.

### DPF-3 is broadly expressed and accumulates in P-granules

The data thus far support a function of DPF-3 in the germline. To visualize *dpf-3* expression, we inserted a sequence encoding a 3xFLAG::TEV::GFP::HiBit tag just before the stop codon of the endogenous *dpf-3* gene. The resulting *dpf-3(xe246)* animals were fertile at both 20°C and 26.5°C, confirming functionality of the tagged protein (fig. S1E). Imaging revealed *dpf-3* expression in various tissues (fig. S2A-D), with particularly high levels in sperm progenitor cells (Fig. 2A) and spermatozoa (Fig. 2B). Moreover, although DPF-3 appeared distributed diffusely in the cytoplasm in most cells, it accumulated in perinuclear punctae in sperm progenitor cells and in the proximal gonad particularly in maturing oocytes (Fig. 2B). Using PGL-1::mTagRFP-T to visualize P-granules (Kawasaki *et al*., 1998), we identified the punctae in maturing oocytes as P-granules, but note that not all P-granules also exhibited DPF-3 accumulation (Fig. 2C; fig. S2E). P-granules are known to accumulate several proteins important for small RNA mediated gene regulation (Seydoux, 2018), and we explore and confirm the hypothesis that DPF-3 also function in this process in a later section.

**Fig. 2.**
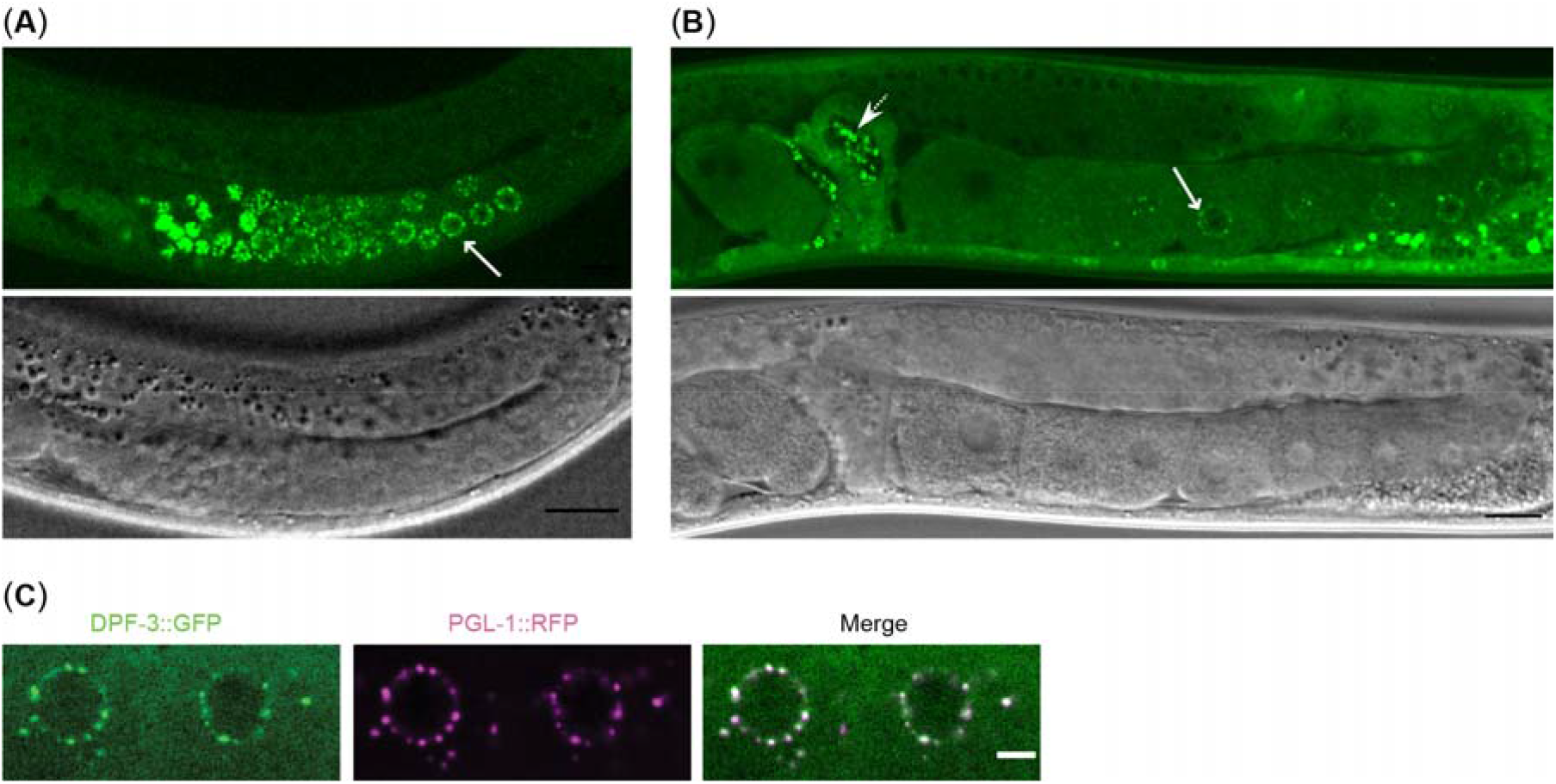
DPF-3 accumulates prominently in the germline. (**A-B**) Single plane confocal images showing the *dpf-3* expression in the gonad of L4 stage (A) and adult (B) animals. White arrows indicate punctate signal in sperm progenitor cells (A) and maturing oocytes (B). A dashed white arrow indicates spermatids (B), scale bar is 10 μm. (**C**) Representative images of developing oocytes from a strain expressing *dpf-3::gfp* (green) and *pgl-1::mTagRfp-t* (purple). Merge shows the co-localization, scale bar is 5 μm.

### Loss of DPF-3 causes TE de-repression, DNA damage and cell death

To understand the underlying cause of the temperature-dependent fertility defect and identify differentially expressed genes, we performed total RNA sequencing. We included *mut-2(ne3370)* and *mut-7(ne4255)* mutant animals in these experiments, which also exhibit temperature-dependent fertility defects (Chen *et al*., 2005; Ketting *et al*., 1999). As hermaphrodite sperm production is restricted to the last (L4) larval stage, we examined populations of synchronized L4 stage animals. To minimize secondary effects,we performed the experiment at 15°C, a temperature where all the *dpf-3* mutant animals examined produce sperm and are fully fertile.

We observed significant transcript dysregulation under these conditions in *dpf-3Δ, mut-2* and *mut-7* mutant relative to wild-type animals (Fig. 3A), with de-silencing of several Transposable Elements (TEs) (Fig. 3B and fig. S3A). Even when increases appear modest at the aggregated family levels, individual family members exhibit substantial transcript accumulation, as illustrated by *ZC15.3*, which harbors a pao class retrotransposon element belonging to the CER12 family (Fig. 3C, D and fig. S3B). Hence, loss of *dpf-3* appears to desilence specific TEs, while *mut-2* or *mut-7* loss leads to a broader defect (Fig. 3A and fig. S3A).

**Fig. 3.**
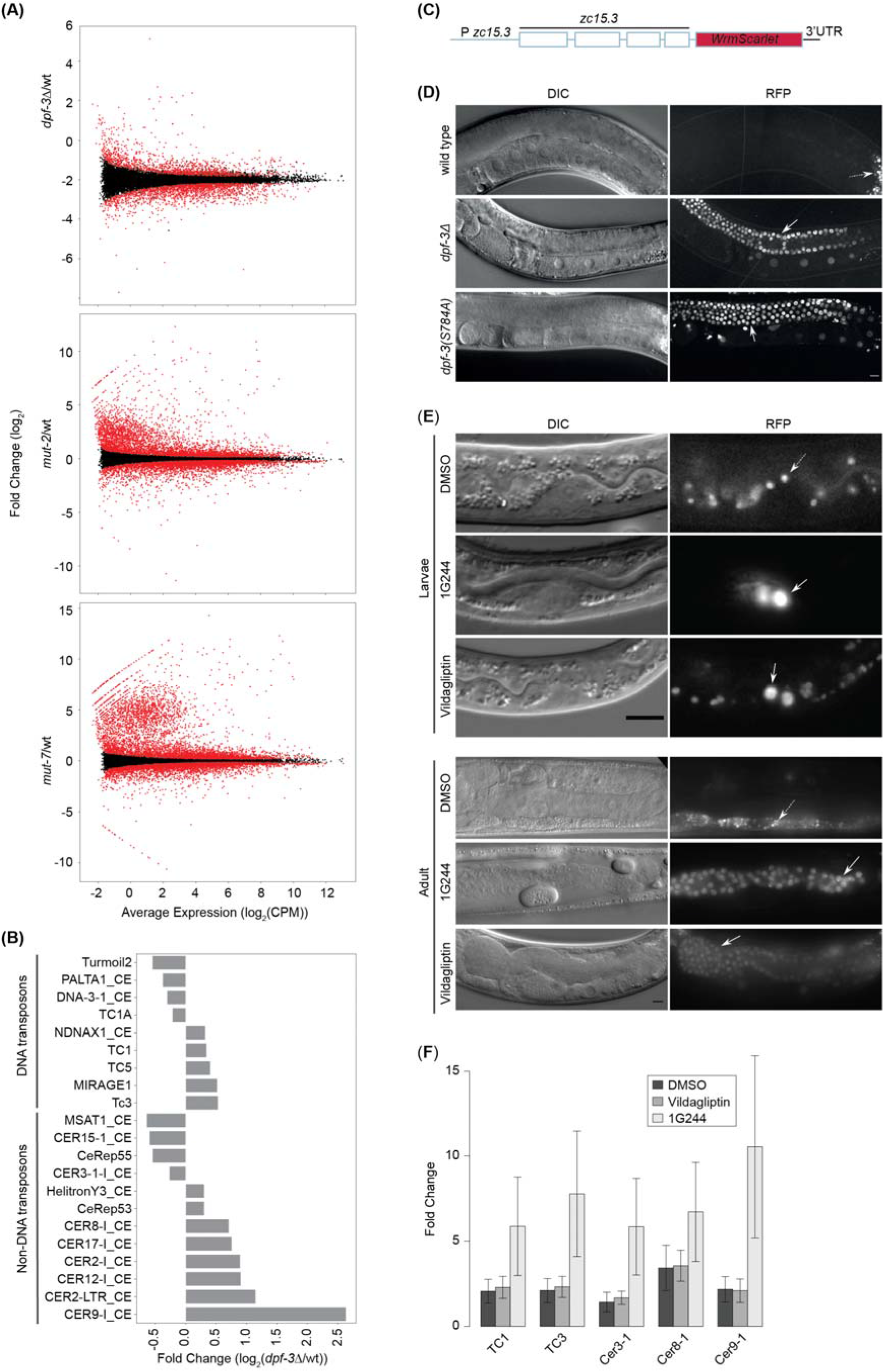
Loss of *dpf-3* leads to transposon de-silencing and DNA damage. (**A**) MA plot showing the differential expression (fold change in log2) of transcripts in the indicated mutant relative to wild-type animals plotted over average expression in the respective mutant and wild-type condition (as CPM – Counts Per Million mapped reads). Each dot represents an individual transcript; significantly changing (FDR<0.05) transcripts are colored red. (**B**) Barplot of DNA and non-DNA transposons at the repeat level that are significantly changed in *dpf-3Δ* mutant compared to wild-type animals. The repeat names obtained from the UCSC genome browser are shown. The x-axis indicates the fold change. (**C**) The architecture of the *ZC15.3*::wrmScarlet transgene reporter. (**D**) Single plane DIC and fluorescent confocal images of wild-type and *dpf-3* mutant adult animals expressing *ZC15.3*::wrmScarlet transgene grown at 15°C. Scale bar is 10 μm. (**E**) DIC and fluorescent images of larvae (above) and adult (below) animals expressing the *ZC15.3*::wrmScarlet transgene grown at 26.5°C treated with vehicle control (DMSO) or DPP8/9 inhibitor 1G244 at 20 μM or DPPIV inhibitor vildagliptin at 200 μM concentration. The animals were treated with drug/control from embryo stage. Scale bar is 10 μM. Dashed white arrow indicates autofluorescence, white arrow germline progenitor cells (Larvae, top panel) and germline (Adult, bottom panel). (**F**) RT-qPCR quantification of the indicated transposons. The average of three biological replicates is shown. Error bars indicate Standard Error of Mean (SEM). The RNA was extracted from animals expressing the *ZC15.3*::wrmScarlet transgene grown at 26.5°C and treated as indicated.

To visualize up-regulation of TE-containing transcripts, we fused *ZC15.3* to a sequence encoding wrmScarlet (red) fluorescent protein (Fig. 3C). We observed a dramatic increase in the levels of the fusion protein in *dpf-3* mutant compared to wildtype animals specifically in the gonad. De-silencing occurred in all developmental stages that we examined (fig. S3B), including adulthood (Fig. 3D), when only oocytes are generated.

Unprogrammed repeat and transposon expression is known to result, independent of transposition activity, in DNA damage and cell death in the *C. elegans* germline (Zeller *et al*., 2016), providing a potential explanation for the lack of sperm progenitor cells and mature sperm that we had observed in *dpf-3* mutant animals (Fig. 1D). Indeed, when we examined the gonads of *dpf-3* mutant animals for DNA damage by anti-RAD-51 antibody staining, we observed a greatly increased number of RAD-51 punctae in the mutant relative to wild-type gonads (fig. S3C-E). Collectively, these data support a scenario where de-silencing of TEs causes, or contributes to, DNA damage and sterility of *dpf-3* mutant animals.

### Inhibition of DPF-3 causes an acute TE silencing defect

To explore the kinetics of TE desilencing and confirm that it depended on DPF-3 enzymatic activity, we turned to using chemical inhibitors. We exposed animals carrying the *ZC15.3*:: wrmScarlet transgene to 1G244, a specific inhibitor of the DPF-3 orthologues DPP8/9 (Wu *et al*., 2009), or vildagliptin, which, at low doses inhibits DPPIV but at high doses also inhibits DPP8/9 (Villhauer *et al*., 2003; Burkey *et al*., 2008). Indeed, recapitulating the *dpf-3* mutant phenotype, when grown at 26.5°C, 70% (52/75) of P0 animals (see methods) exposed to 20 μM IG244 exhibited TE reporter desilencing (Fig. 3E). A ten-fold higher concentration (200 μM) of vildagliptin was required to observe a comparable effect (80% (50/63 animals) wrmScarlet positive). RT-qPCR confirmed desilencing of additional RNA and DNA transposons on bulk animals (Fig. 3F). Strikingly, desilencing occurred rapidly, within one generation, distinguishing it from the chronic effects that occur upon long-term loss of other TE silencing factors (Sakaguchi *et al*., 2014; Tabara *et al*., 1999)

### A subset of 22G RNAs is depleted in *dpf-3* mutant animals

Although the temperature-sensitive sterility and TE silencing defects in *dpf-3* mutant animals appeared consistent with impaired small RNA activity, the rapid onset was unexpected. Hence, we used a 5’-independent cloning protocol to quantify small RNAs from wild-type and mutant animals of L4 stage grown at 15°C. While little or no change occurred for 21U RNAs (Fig. 4A and fig. S4A), we observed that 22G RNAs, which comprise most of the mapped reads (fig. S4C), were significantly affected by the *dpf-3* status: The levels of a substantial number of 22G RNAs decreased in *dpf-3* mutant animals, and this effect was comparable qualitatively and quantitatively between *dpf-3Δ* and *dpf-3(S784A)* mutant animals (Fig. 4B). Again, *mut-2* (Fig. 4B) and *mut-7* (fig. S4B) mutant animals exhibited an overlapping, yet broader defect in 22G RNA accumulation. Notably, the transcripts that were strongly derepressed in *dpf-3Δ* mutant animals also showed decreased levels of 22G RNAs coming from the anti-sense strand of the respective genomic region (Fig. 4C, D). As in *mut-2* mutant animals, however, not all transcripts with decreased 22G RNA targeting were upregulated, consistent with redundant silencing mechanisms acting on them (Padeken *et al*., 2019). We conclude that DPF-3 is required for accumulation of a subset of endo-siRNAs that normally silence transposons and repeats.

**Fig. 4.**
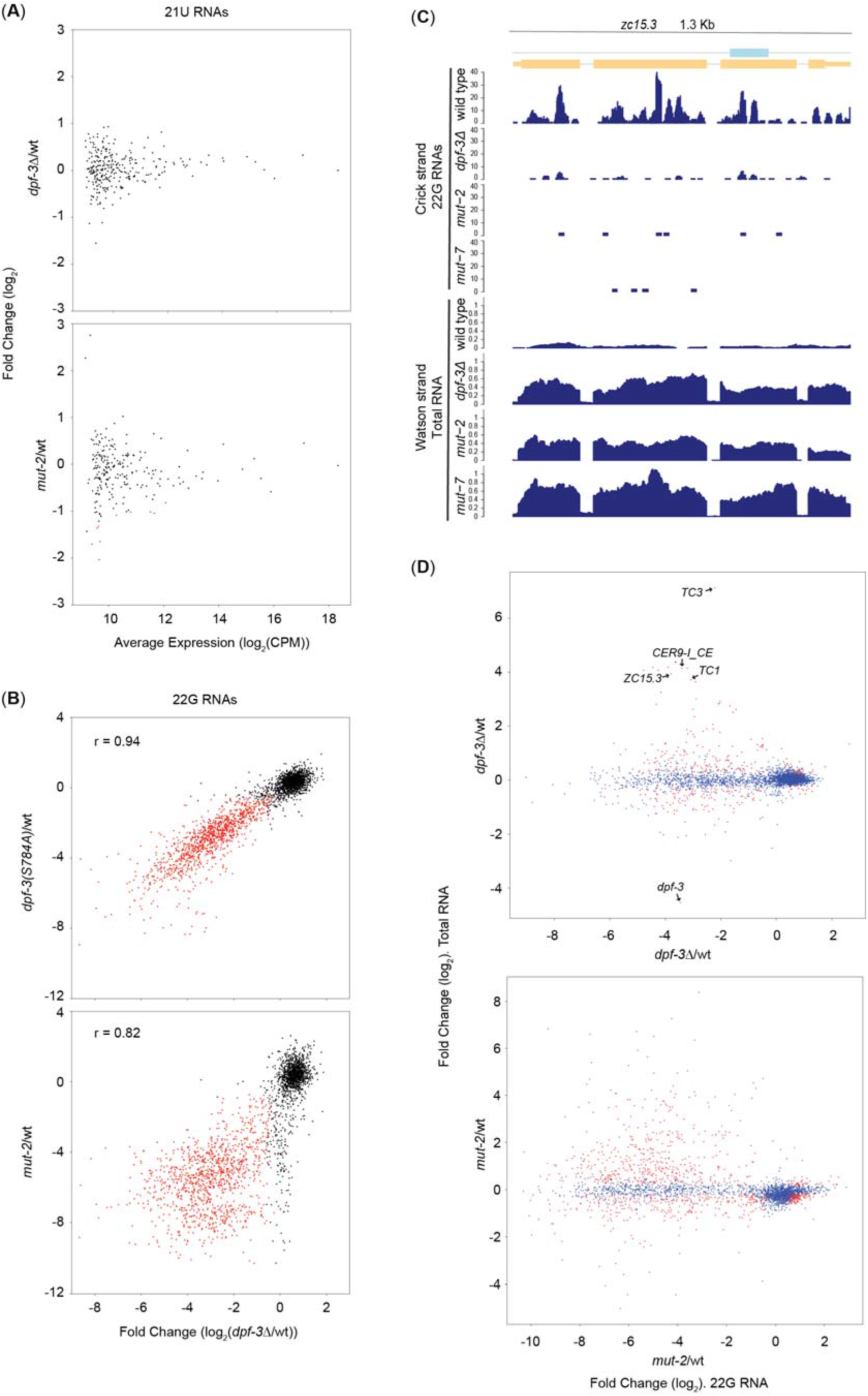
Specific 22G RNAs are down-regulated in *dpf-3* mutant animals. (**A, B**) Scatter plots showing differential expression of 21U RNAs (A) or 22G RNAs (B) between wild-type and the indicated mutant animals. Each dot represents an individual 21U RNA (A) or all the 22G RNAs that match in anti-sense orientation to a given transcript (B); red indicates a significant depletion in both mutant backgrounds (FDR<0.05). The r values indicate the Pearson correlation coefficient. (**C**). Genomic track showing the *ZC15.3* locus transcribed from Watson strand. The pao LTR retrotransposon element present in the *ZC15.3* is indicated by a blue rectangle. Note that 22G RNAs that are antisense to *ZC15.3* are present in wild-type and depleted in all the mutant animals tested. Contrarily, the ZC15.3 transcript is significantly upregulated in mutant but not in wild-type animals. (**D**) Scatter plot showing the differential expression of total RNA over 22G RNAs in *dpf-3Δ* (above) and *mut-2* (below) mutant relative to wild-type animals. Each dot represents the combined data of total and small RNA sequencing data: a sense transcript (total RNA, y-axis) and all the 22G RNAs that match in anti-sense orientation to that particular transcript (x-axis). Red color indicates a significant change (FDR <0.05) in both total and 22G RNA levels. Selected up-regulated TEs at the individual transcript level in *dpf-3Δ* are indicated. Gene identities obtained from wormbase.org.

### 22G RNAs lost in *dpf-3* mutant animals are loaded into WAGOs

22G RNAs are bound by, and function with Ago, more specifically, WAGOs and CSR-1 proteins (Youngman and Claycomb, 2014). Whereas WAGOs repress their targets including TEs (Gu *et al*., 2009; Yigit *et al*., 2006), CSR-1 promotes the expression of endogenous germline genes (Claycomb *et al*., 2009). Examination of Ago protein N-terminal sequences revealed for WAGO-1, WAGO-3 and WAGO-4 either alanine or proline in position two and/or three, qualifying them as potential DPF-3 substrates. We wondered whether DPF-3 might affect 22G RNAs accumulation through altering the levels or functionality of these WAGOs, and, hence, sought to define their 22G RNA cargo. Indeed, when we immunoprecipitated WAGO-1 or WAGO-3 from L4-stage wildtype animals grown at 15°C and sequenced the bound small RNAs (IP-seq), we observed a large overlap of 22G RNAs that are bound by either WAGO and lost in *dpf-3* or *mut-2* mutant animals. By contrast, for CSR-1, only a marginal overlap occurred between its associated 22G RNAs and the 22G RNAs that are lost in *dpf-3* or *mut-2* mutant animals (Fig. 5A and fig. S5A).

**Fig. 5.**
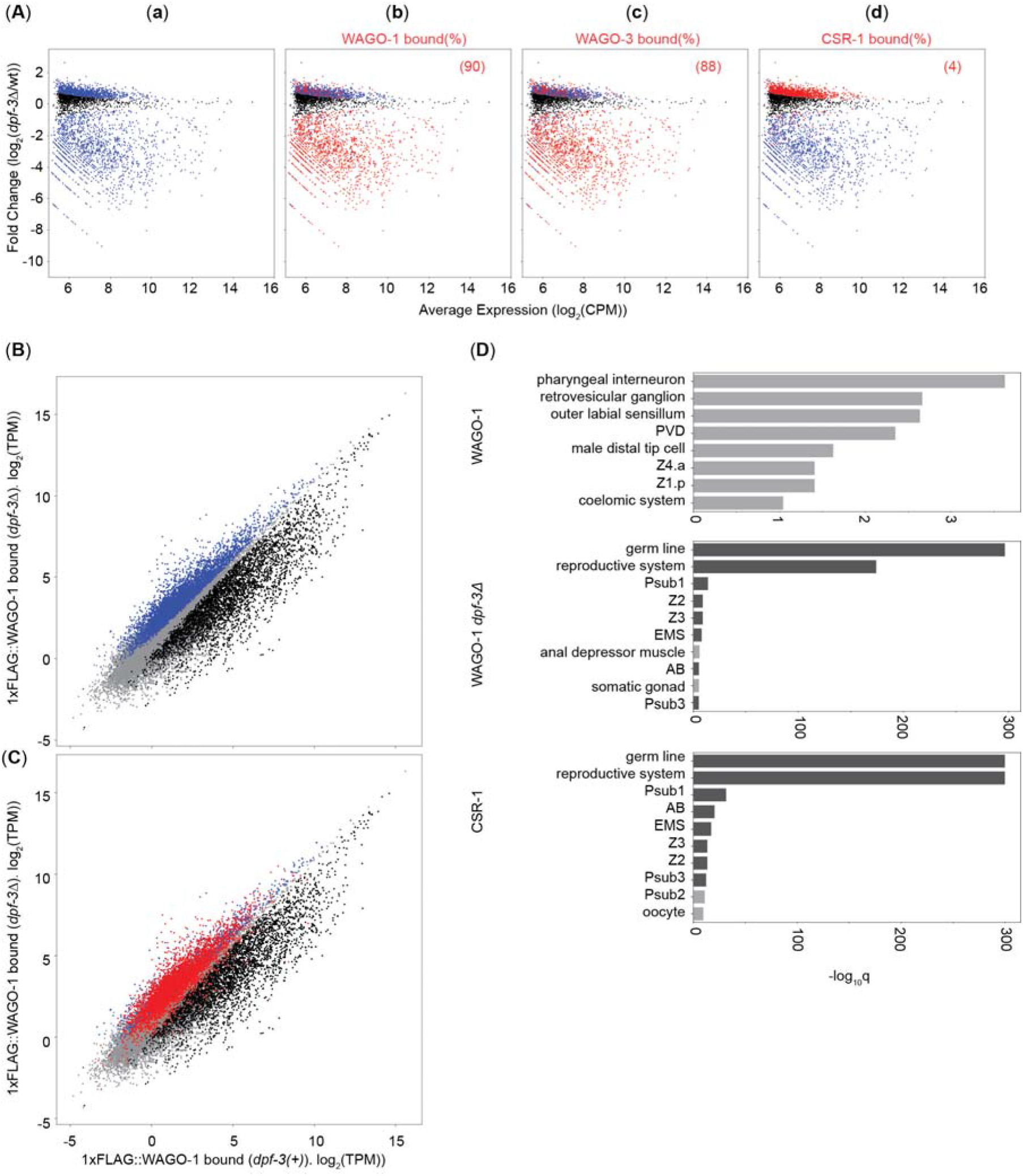
Loss of *dpf-3* alters the cellular and WAGO-1-bound 22G RNA complement. (**A**). MA plots showing the differential expression of 22G RNAs in the *dpf-3Δ* mutant relative to wild-type animals plotted over average expression in the respective mutant and wild-type condition (as CPM – Counts Per Million mapped reads). Each dot represents all the 22G RNAs that match in anti-sense orientation to a given transcript. Significantly changing (FDR<0.05) transcripts are colored blue. In panels (**b**-**d**), differentially expressed 22G RNAs that are bound by a specific Ago protein are colored red. The number (%) shown in each individual plot indicates the fraction of 22G RNAs bound by a specific Argonaute protein and down regulated in *dpf-3Δ* mutant animals (e.g. 90% of 22G RNAs bound by WAGO-1 are down-regulated in *dpf-3Δ* mutant animals (b)). (**B**) Scatter plots showing binding of 22G RNAs to WAGO-1 in lysates from *dpf-3Δ* (y-axis) and wild-type (x-axis) L4 stage animals. Each dot represents all the 22G RNAs that match in anti-sense orientation to a given transcript. Gray indicates no change, black significant depletion and blue significant enrichment on WAGO-1 *in dpf-3Δ* mutant relative to wild-type animals (FDR<0.05). (**C**) Scatter plot exactly as shown in (B), but the 22G RNAs that are bound by CSR-1 are indicated in red color. 63% (3384/5359) of 22G RNAs bound to WAGO-1 in *dpf-3Δ* background are also bound by CSR-1, while only 3% (134 out of 3973) of 22G RNAs bound to WAGO-1 in wild-type condition are bound by CSR-1. (**D**) Tissue Enrichment Analysis (TEA) of 22G RNAs bound to WAGO-1 in wild-type (black points from (B)) or in *dpf-3Δ* mutant animals (blue points from (B)), or bound by CSR-1. Tissues that are enriched in both the last two comparisons are colored dark grey. The x-axis indicates *q* values (-log_10_).

### DPF-3 prevents accumulation of small RNAs targeting germline genes on WAGO-1

We wondered whether in addition to promoting accumulation of *bona fide* 22G RNAs on WAGO proteins, DPF-3 might also prevent binding of other small RNAs, and thus potentially protecting the host from detrimental silencing of self genes. IP-seq of WAGO-1 from either wild-type or *dpf-3Δ* mutant animal lysate revealed the enrichment of “neo-22G RNAs” in the mutant background (Fig. 5B), which overlapped extensively with CSR-1-bound 22G RNAs (Fig. 5C). Substantiating this observation further, the “neo-22G RNAs” bound to WAGO-1 in *dpf-3Δ* mutant animals are significantly enriched for germline-expressed genes, distinguishing them from the canonical WAGO-1-associated 22G RNAs in wild-type animals (Fig. 5D). Hence, in the absence of DPF-3, WAGO-1 acquires non-canonical 22G RNAs targeting germ line genes, which may contribute to the sterility phenotype.

### DPF-3 is an N-terminal dipeptidase that processes WAGO-1 and WAGO-3 *in vivo*

To examine whether DPF-3 has dipeptidase activity acting on WAGOs, we incubated immunoprecipitates from *dpf-3(+)* and *dpf-3(S784A)* animals with peptide substrates representing the N-termini of WAGO-1 and WAGO-3 and quantified changes in the predicted product, shortened by two amino acids, by mass spectrometry over time. We observed substrate cleavage, and generation of the expected product in the presence of wild-type DPF-3 protein (Fig. 6A). By contrast, *DPF-3(S784A)* mutant protein, although immunoprecipitated at similar levels (fig. S6A), failed to generate product. This confirms that DPF-3 is an N-terminal dipeptidase, which is rendered inactive by the S784A mutation *in vitro*.

**Fig. 6.**
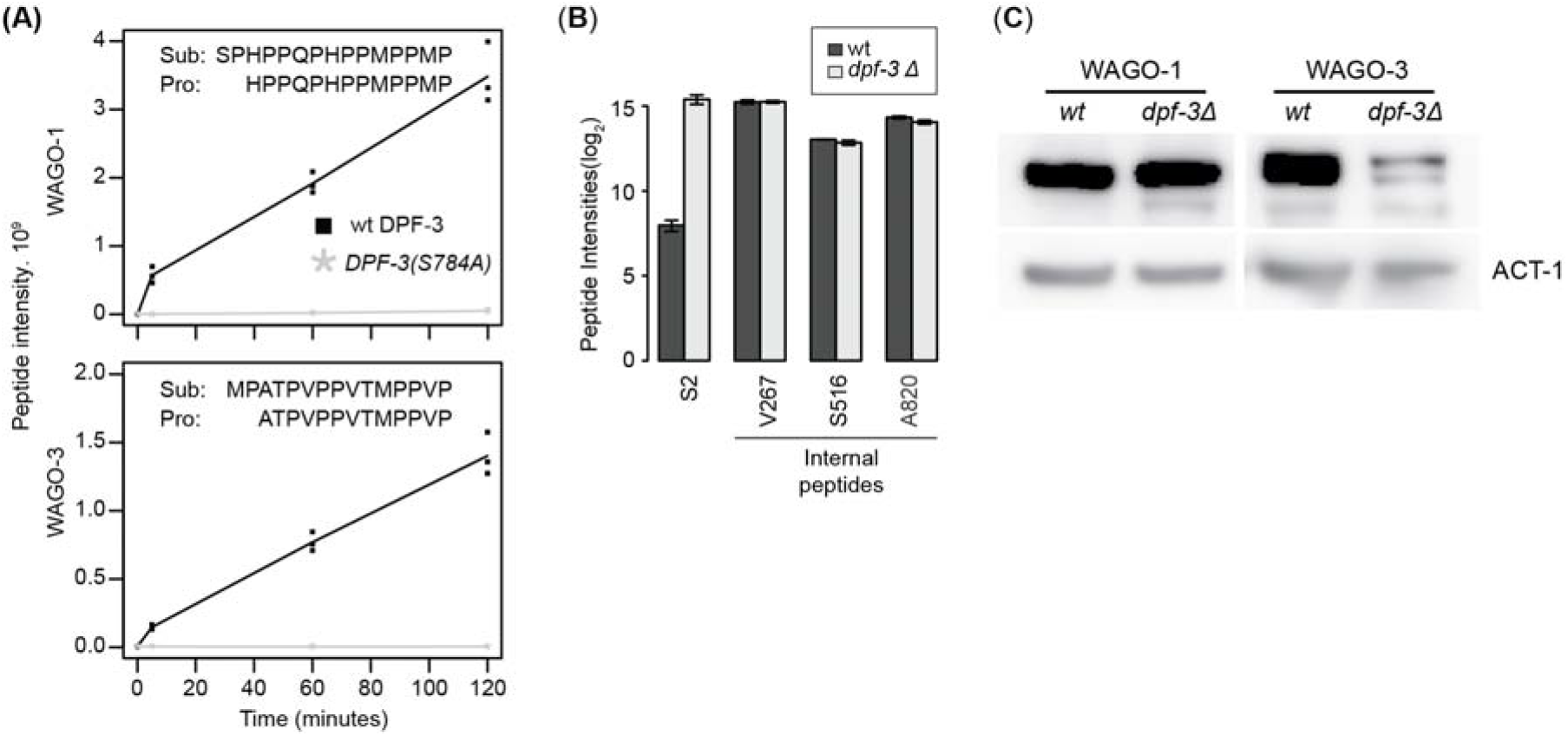
WAGO-1 and WAGO-3 are processed by DPF-3. (**A**) Product peptide intensity of WAGO-1 (top) and WAGO-3 (bottom) plotted over reaction time. Data points of three biological replicates are shown; a line indicates mean values. Since no product was seen when *DPF-3(S784A)* mutant protein was used, all the relevant replicate points are on top of each other. The sequences of the substrates (Sub) and the products (Pro) are shown. (**B**) Bar plot of different peptides (x-axis) showing average peptide intensity (y-axis, log_2_) from three biological replicates. For details about the different peptides labelled on the x-axis, see “peptide description” in materials and methods section. Error bars represent Standard Error of Mean (SEM). (**C**) Western blot probed with anti-FLAG Antibodies to detect the indicated, endogenously FLAG-tagged, WAGO proteins in wt or *dpf-3Δ* mutant animals. ACT-1 serves as a loading control.

To confirm that DPF-3 processes WAGO-1 and WAGO-3 *in vivo*, we used massspectrometry to map the N-termini of the endogenous, FLAG-tagged proteins, immunopurified from either wild-type or *dpf-3Δ* mutant animals. We inserted the FLAG tags between amino acids 45 and 46 for WAGO-1 and 46 and 47 for WAGO-3 to avoid interference with N-terminal processing. Whereas “internal peptides” away from the N-terminus were similarly abundant in wild-type and *dpf-3* mutant immunoprecipitates, specific differences occurred for peptides derived from the N-terminus. Thus, a fulllength N-terminal peptide (starting at S2, due to de-methionylation) of WAGO-1 was ~170 fold more abundant in *dpf-3Δ* than wild-type animals (Fig. 6B). Moreover, although the levels of the predicted product (starting with H4) were comparable between the conditions, several more extensively truncated N-terminal WAGO-1 peptides accumulated preferentially in the presence of DPF-3 (fig. S6C, D; particularly pronounced for peptides starting at P16, V18 and P29), suggesting further processing after the initiating cleavage. The proteolytic processing did not appear to affect WAGO-1 stability, as steady-state levels were comparable in wild-type and *dpf-3* mutant lysates (Fig. 6C and fig. S6B).

As for WAGO-1, we observed enhanced accumulation of N-terminally truncated peptides of WAGO-3 in wild-type relative to *dpf-3Δ* mutant animals (fig. S6C and E). However, the steady state levels of WAGO-3 significantly decreased in *dpf-3Δ* mutant animals (Fig. 6C and fig. S6B). Hence, proteolytic activity of DPF-3 on WAGO-3 appears to promote rather than antagonize its stability and activity.

### WAGO-1 and WAGO-3 are critical substrates of DPF-3

To test whether and to what extent processing of WAGO-1 and WAGO-3 could explain DPF-3’s physiological function, we mutated the endogenous WAGO-1 and WAGO-3 proteins to render them refractory to DPF-3-mediated cleavage. While point mutations in individual *wago* genes did not show any significant phenotype, the *wago-1(P3G), wago-3(P2G,A3E/R)* double mutant animals extensively recapitulated *dpf-3* mutant phenotypes: First, they had greatly reduced brood sizes at 25°C (Fig. 7A), and they were 100% sterile at 26.5°C (n>100). Second, the WAGO-3 protein levels decreased (Fig. 7B and fig. S7A). Third, the *ZC15.3*::wrmScarlet TE reporter transgene was de-silenced (fig. S7B). Fourth, a fraction of 22G RNAs was depleted, and this fraction greatly overlapped with the subset of 22G RNAs depleted in *dpf-3* mutant animals (Fig. 7C). Finally, immunoprecipitation followed by small RNA sequencing of WAGO-1 from *wago-1(P3G), wago-3(P2G,A3E)* double mutant animal lysate revealed the enrichment of “neo-22G RNAs” (Fig. 7D), similar to *dpf-3Δ* mutant animals (Fig. 5B). These “neo 22G RNAs” also overlapped extensively with CSR-1-bound 22G RNAs (Fig. 7E) and they were significantly enriched for targeting germline expressed genes (fig. S7C). We conclude that WAGO-1 and WAGO-3 are the critical substrates of DPF-3, that mediate the function of this protease in fertility and TE silencing.

**Fig. 7.**
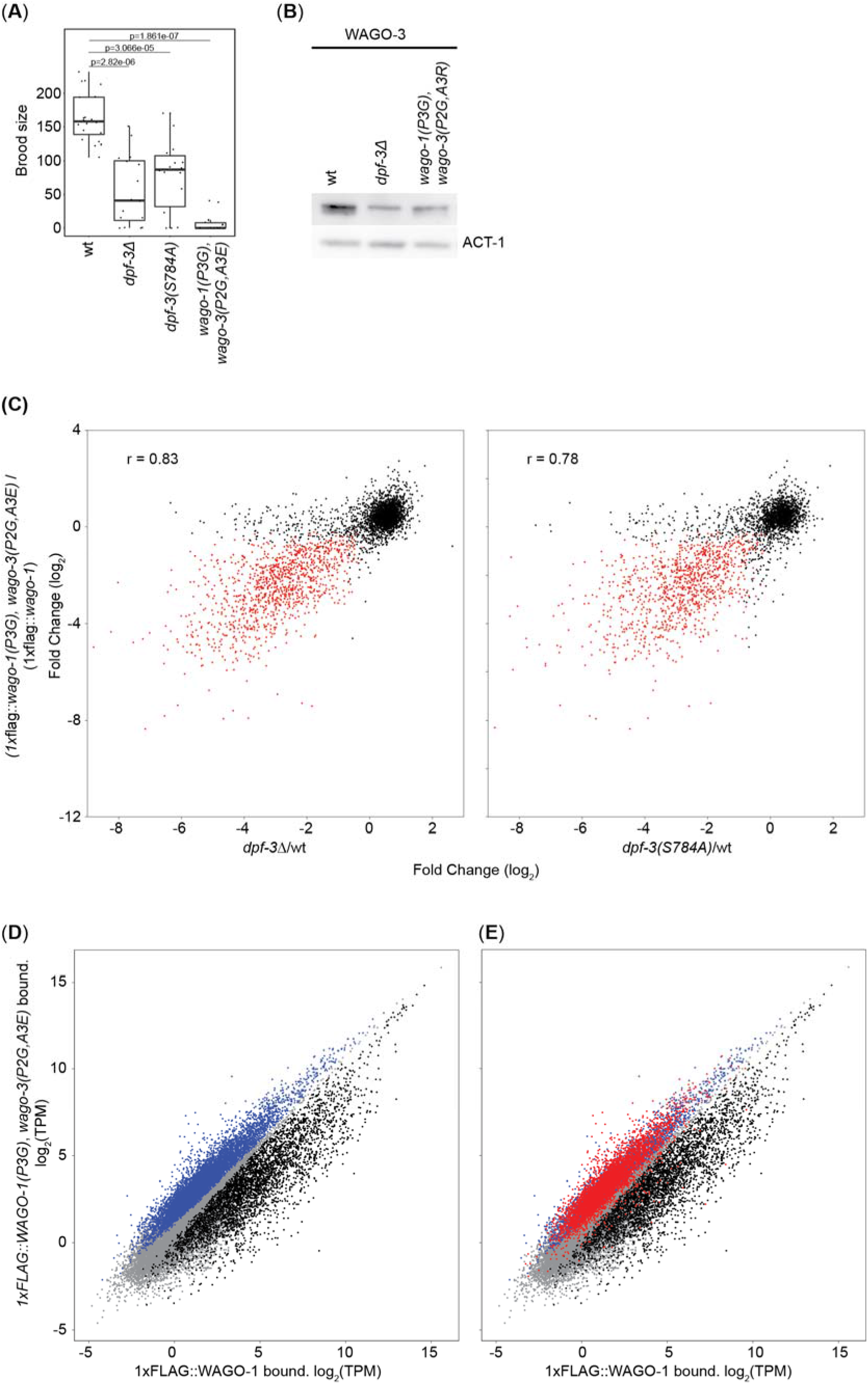
Animals producing DPF-3-refractory WAGO-1 and WAGO-3 exhibit *dpf-3* loss-of-function phenotypes. (**A**) Box plot of the brood sizes of the indicated strains, grown at 25°C. Each dot represents an individual worm. Wild-type and *dpf-3* mutant animal data are from Fig. 1B and included for comparison. p values calculated using Wilcoxon test is shown. (**B**) Western blot probed with anti-FLAG antibodies to detect the endogenously FLAG-tagged WAGO-3 protein in wt or indicated mutant animals. ACT-1 serves as a loading control. (**C**) Scatter plots showing differential expression of 22G RNAs between wild-type and the indicated mutant animals. Each dot represents all the 22G RNAs that match in anti-sense orientation to a given transcript; red indicates a significant depletion in both mutant backgrounds (FDR<0.05). The r values indicate the Pearson correlation coefficient. (**D**) Scatter plots showing binding of 22G RNAs to WAGO-1 in lysates from *wago-1(P3G), wago-3(P2G,A3E)* mutant and wild-type L4 stage animals. Each dot represents all the 22G RNAs that match in anti-sense orientation to a given transcript. Gray indicates no change, black significant change depletion and blue significant enrichment on WAGO-1 in *wago-1(P3G), wago-3(P2G,A3E)* mutant relative to wild-type animals (FDR<0.05). (**E**) Scatter plot exactly as shown in (D), but the 22G RNAs that are also bound by CSR-1 are indicated in red color. 54% (4054/7406) of 22G RNAs bound to WAGO-1 in *wago-1(P3G), wago-3(P2G,A3E)* background are also bound by CSR-1, while only 2% (87/4026) of 22G RNAs bound to WAGO-1 in wild-type condition are bound by CSR-1.

## Discussion

The heritable nature of small RNA-mediated gene silencing is both a strength and a liability of this defense system against selfish genetic elements. It provides a potent mechanism to safeguard genome integrity in the germline, yet risks collateral damage if not carefully constrained. Our work identifies the DPF-3 protease, and specifically the proteolytic processing of WAGO-1 and WAGO-3 that it mediates, as a mechanism that promotes WAGO association with non-self targeting 22G RNAs over self targeting, CSR-1-type, 22G RNAs.

The same species of RNA, 22G RNAs, has two opposing functions in the *C. elegans* germline, namely promotion and silencing of gene expression, depending on the identity of the Ago protein with which it associates. Hence, 22G RNA binding to, and/or retention on, WAGOs versus CSR-1 must be tightly controlled. Indeed, discrimination appears normally robust, and even in the absence of factors promoting WAGO-type 22G RNA accumulation such as PRG-1 and ENRI-3, 22G composition drifts rather slowly such that fertility declines gradually, over several generations (termed a mortal germline (Mrt) phenotype) (Lewis *et al*., 2020; Barucci *et al*., 2020; Spracklin *et al*., 2017). Yet how such a discrimination is achieved, has remained largely unknown.

Our results have revealed an essential role of proteolytic, N-terminal processing of WAGO-1 and WAGO-3 in this process: Inactivation of DPF-3, which normally mediates this processing, depletes the normal complement of 22G RNAs on WAGO-1, promotes misloading with CSR-1-type 22G RNAs and destabilizes WAGO-3. TE desilencing, DNA damage, and acute sterility ensue. Loss of male fertility, and specifically absence of sperm, is a major cause of sterility in *dpf-3* mutant animals, but TE upregulation occurs across the gonad, including at the adult stage, when sperm is no longer produced, and wild-type sperm can only partially restore fertility of *dpf-3* mutant hermaphrodites. Hence, DPF-3-mediated processing of WAGO-1 and WAGO-3 appears important for both male and female fertility, functioning in both the male and the female germline.

*In vitro* processing assays and examination of WAGO-1 and WAGO-3 N-termini *in vivo* identify WAGO-1 and WAGO-3 as DPF-3 substrates. We expect that DPF-3 may process additional substrates, yet, the two Ago proteins are clearly key mediators of its function in fertility: Mutations that render them resistant to processing by DPF-3 recapitulate *dpf-3* loss of function phenotypes extensively, extending, most notably, to comparable changes in 22G RNA levels, at the level of both total cellular small RNA and WAGO-1-bound small RNAs.

Strikingly, the onset of phenotypes seen upon *dpf-3* mutation or chemical inhibition is rapid, with TE desilencing and sterility occurring within one generation. Also unlike in *prg-1* mutant animals (Barucci *et al*., 2020), there is no evidence of specific mis-loading of 22G RNAs anti-sense to histone protein coding genes into WAGO-1 (fig. S5B) nor of decreased levels of histone protein coding mRNAs (fig. S5C). Hence, rather than maintaining over time the composition of the 22G RNA complement associated with WAGOs, DPF-3 appears to play a major role in generating such a distinct pool of WAGO-bound 22G RNAs, possibly by preventing an erroneous, stochastic association of various types of 22G RNAs with WAGO-1.

Ago proteins appear particularly diverse at their N-termini, implying a potential for regulatory functions. Further supporting this notion, the N-terminal domain of human Ago1 blocks the protein’s catalytic activity in RNA cleavage (Hauptmann *et al*., 2013), and specific mutations in the N-terminal domain (NTD) of human AGO2 modulate its ability to complex with small RNAs (Kwak and Tomari, 2012). While it remains to be established that these features are naturally exploited for regulation, we are particularly intrigued by the observation that AGO2 NTD mutations can confer specificity for particular RNA substrates (Kwak and Tomari, 2012): whereas binding of siRNAs, derived from fully complementary duplexes, is blocked, binding of miRNAs, derived from partially complementary duplexes, proceeds normally. Although synthesis of 22G RNAs through RNA-dependent RNA polymerase (RdRPs) can be assumed to involve a fully complementary intermediate, it is not known how loading of WAGOs or CSR-1 occurs, and, with two distinct RdRPs synthesizing 22G RNAs, i.e., RRF-1 the WAGO-type (Gent *et al*., 2010), EGO-1 the CSR-1 type (Claycomb *et al*., 2009), this may indeed also differ between these classes.

The separation of 22G RNA production by RdRP provides an additional opportunity for regulation, based on spatial separation: whereas RRF-1 accumulates in mutator foci (Phillips *et al*., 2012), EGO-1 localizes preferentially to P-granules (Claycomb *et al*., 2009). Yet, unexpectedly, CSR-1, WAGO-1, and WAGO-3 all accumulate in P-granules (Gu *et al*., 2009; Claycomb *et al*., 2009) (J.S. and R.K., unpublished data), as does DPF-3. Although further work is needed to confirm that the sites of steady-state localization of these different proteins are also their sites of action, these findings imply that WAGO-1 and WAGO-3 and/or the 22G RNAs destined to bind them are able to move from mutator foci to P-granules. Speculatively, DPF-3-mediated processing might affect such dynamics, e.g., by promoting WAGO shuttling through mutator foci.

If we accept that 22G RNA loading onto WAGOs occurs in mutator foci, but that DPF-3 processes WAGOs in P-granules, where both accumulate, we may also speculate that N-terminal WAGO processing prevent 22G RNA release and reloading rather than promote loading itself. In this view, a more “open” conformation of N-terminally extended WAGOs would promote both binding and unbinding of 22G RNAs, while its removal would “clamp” the WAGO on a loaded RNA, preventing it from exchanging its proper 22G RNA cargo with a CSR-1-type 22G RNA in the hostile environment of a P-granule.

Finally, we note that an effect on RNA binding may also explain the destabilization of WAGO-3 that we observe in *dpf-3* mutant animals: although this could be a direct consequence of DPF-3-mediated processing normally inactivating an N-terminal degron on WAGO-3, it might alternatively result from decreased stability due to impaired loading with small RNAs, as previously reported for other Agos (Smibert *et al*., 2013). Irrespective of which explanation applies, we note that the effect of DPF-3-mediated processing on WAGO-1 and WAGO-3 differs from that of other DPPIV family proteases on their substrates (Mentlein, Gallwitz and Schmidt, 1993; Justa-Schuch *et al*., 2016): instead of increasing catabolism, we observe functional modification, illustrated by the change of 22G RNAs on WAGO-1. We propose that these may be relevant outcomes for other members and substrates of the DPF proteins, which should be considered in understanding the physiological functions of this biomedically important family of proteases.

## Acknowledgements

We thank Anca Neagu, Lan Xu and Iskra Katic for technical support, Laurent Gelman and Jan Eglinger for help with microscopy and co-localization analysis, and Kirsten Jacobeit, Eliza Mereno Pandini Figueiredo and Sebastien Smallwood for sequencing. We are grateful to Dimos Gaidatzis, Michael Stadler and Hans-Rudolf Hotz for bioinformatics and to Jan Seebacher and Vytautas Iesmantavicius for mass spectrometry advice and discussion. We thank Thomas Welte for fruitful discussions and Benjamin Towbin, Takashi Miki, Marc Buhler for critically reading the manuscript. We thank Florian Steiner for providing the *his-70::gfp* strain prior to publication and Germano Cecere for providing the *csr-1::3xFLAG::1xHA* strain. Some strains were provided by the *Caenorhabditis* Genetics Center (CGC), which is funded by NIH Office of Research Infrastructure Programs (P40 OD010440).

## Funding

The project was supported through funding from the Swiss National Science Foundation through the National Center of Competence in Research RNA and Disease and the Novartis Research Foundation through the Friedrich Miescher Institute for Biomedical Research (to H.G.). and from the Deutsche Forschungsgemeinschaft (KE 1888/1-1, KE 1888/1-2 and KE 1888/6-1; to R.F.K.).

## Author Contributions

R.K.G. and H.G. conceived the project, designed and analyzed the experiments and wrote the manuscript. H.G. supervised the project. R.K.G. and K.B. performed most of the experiments. F.G. and C.H.S. analyzed the deep sequencing data. D.H. performed and analyzed mass spectrometry. J.S. and R.F.K. provided strains, shared unpublished data and commented on the manuscript.

## Competing interests

The authors declare no competing interests

## Data and material availability

All the data is included in the manuscript or in the supplementary material. The sequencing data is deposited under super series accession number GSE151717at GEO (https://www.ncbi.nlm.nih.gov/geo/query/acc.cgi?acc=GSE151717). A reviewer token for data access is available through the editorial office. Published research reagents from the FMI are shared with the academic community under a Material Transfer Agreement (MTA) having terms and conditions corresponding to those of the UBMTA (Uniform Biological Material Transfer Agreement).

## Materials and methods

### C. elegans

The strains that are used in this study are described in Supp. Table 1. The wild-type strain we used is Bristol N2. The worms were grown on NGM 2% with *Escherichia coli* OP50 bacteria as a food source unless stated differently. For protein IP experiments, worms were grown at 15°C on peptone rich plates with *Escherichia coli* NA22 bacteria as a food source. To synchronize the worms, we extracted the eggs from gravid adults by bleaching solution (30% (v/v) Sodium hypochlorite, 5% chlorine (Thermo Fisher Scientific; 419550010), 0.75M KOH)). The eggs were left in M9 buffer (42 mM Na_2_HPO_4_, 22 mM KH_2_PO_4_, 86 mM NaCl, 1 mM MgSO_4_) without any food for 12-16 hrs, thus generating synchronized, arrested L1 stage worms. The staged L1 worms were plated on the NGM 2% plate containing OP50 bacteria as a food source. Alternatively (when indicated), staged L1 worms were plated on to peptone rich plates containing NA22 bacteria as a food source.

### RNA extraction, library generation and sequencing

Synchronized worms grown at 15°C at the indicated developmental stages were used for extracting the total RNA as described in (Aeschimann *et al*., 2017). Total RNA library preparation was essentially done as described in (Aeschimann *et al*., 2017). Total RNA-seq libraries were generated using the TruSeq Illumina Total RNA-seq (stranded – high input) according to manufacturer’s protocol. Libraries were sequenced on the NextSeq HIGH-OUT 150 Cycle Paired-end (eg. 2 x 75bp). Preparation of Small RNA libraries (both 5’ dependent and in-dependent) and sequencing was done essentially as described in (Brown *et al*., 2017). Briefly, Small RNA fraction was enriched using a microRNA purification kit (Norgen biotek) with 500ng total RNA as input. The small RNA fraction was treated with 20U of RNA 5’ Polyphosphatase (TAP, Lucigen), in a 50μl reaction at 37°C for 1h. TAP-treated RNA was purified using the Norgen Biotek Single Cell RNA Purification kit and eluted in 10μl. 5μl was used as input for small RNA library preparation using the QIAseq miRNA library kit (Qiagen) according to the manufacturer recommendations. Libraries were sequenced on the Illumina HiSeq 2500 (50cycles single-end run).

### Construction of transgenic lines

All the transgenic worms were generated by single copy integration in Chromosome II or III as described in (Frokjaer-Jensen *et al*., 2012). The fragment of interest is amplified by phusion polymerase (ThermoFisher Scientific) as per the manufacturer instructions and cloned into the vector pCFJ150 by Gibson assembly (Gibson *et al*., 2009). CRISPR-Cas9 was used for genome editing as described in (Katic, Xu and Ciosk, 2015; Arribere *et al*., 2014). Not1 linearized pIK198 is used to clone the sgRNAs by Gibson assembly. A mix containing 50ng/ul pIK155, 100 ng/μl of pIK208, 100 ng/μl of pIK198 with a cloned sgRNA, 20ng/ul of AF-ZF-827 primer, 20ng/ul of single stranded primer as a template for homologous recombination targeting the gene of interest is injected into Bristol N2 worms unless indicated. Transgenic worms were isolated as described in (Arribere *et al*., 2014). In most cases, at least two independent lines were obtained. The transgenic lines were confirmed by PCR and backcrossed at least 3 times. All the sgRNA sequences and primers used for introducing desired changes are listed in Supp. Table 1.

### DPF-3 Rescuing transgene construction

Because *dpf-3* is the second gene in a two-gene operon and because some downstream cistrons in operons are expressed both as polycistronic transcripts and from intercistronic promoters (Miki *et al*., 2016), this transgene contained not only 1.744 kb of operon upstream region as the putative operon promoter, but also the coding region of the first gene, *ahcy-1*, and the operon linker sequence. To prevent overexpression of the first gene in the operon, we mutated the start codon of *ahcy-1*, to prevent its translation, and expressed the transgene in a *dpf-3Δ* mutant background. The plasmid encoding this transgene is made in the following way. PCR with RG630-RG649 and RG650-RG651 and RG652-RG645 using genomic DNA as a template. A gibson assembly reaction was performed as described in (Gibson *et al*., 2009) by mixing these three PCR products and AvrII/BglII linearized pCFJ150 plasmid generating *dpf-3* operon with a mutation of the start codon of the *ahcy-1* gene.

### Image acquisition

All confocal images were acquired exactly as described in (Aeschimann *et al*., 2017). Fluorescent and Differential Interference Contrast (DIC) images represented in Fig. XXX were obtained with Zeiss Axio Observer Z1 microscope and AxioVision SE64 software and Zen 2 (blue edition). Fiji was used for selecting/cropping and processing of the images.

### Construction of Fluorophore tagged *dpf-3* and *pgl-1*

Using the plasmid pIK377 as a template that contains 3xFLAG::TEV::GFP::HiBit, PCR was done with DPF-3 overhang fw-oIK1315 and DPF-3 overhang rev-oIK1288 to generate final asymmetric PCR product containing the intended tag and homology (120 nucleotides) to *dpf-3* gene and tagged lines were obtained as described in (Dokshin *et al*., 2018). The tag in the *dpf-3* is inserted just before the translation stop codon. The RFP tagged *pgl-1* strain is a kind gift from Ketting laboratory and is described in the submitted manuscript (Schreier et al, unpublished).

### Co-localization image analysis

The worms expressing DPF-3::GFP or PGL-1::RFP or both the fluorophores, as well as the fluorescent beads (Molecular probes, Ref-T14792), were imaged in both GFP and RFP channels. Channel correction and co-localization analysis was performed using KNIME Analytics Platform (version 4.1.0, KNIME GmbH). The channel shift for the two-camera acquisition systems was corrected using fluorescence bead images as reference. After channel-shift correction, the GFP- and RFP-positive spots were detected using a Laplacian-of-Gaussian (LoG) detector from Trackmate (Tinevez *et al*., 2017) and an adapter plugin for KNIME (fmi-ij2-plugins-0.2.5https://doi.org/10.5281/zenodo.1173536). To assess co-localization, we measured the correlation of GFP-positive and RFP-positive spots and visualized the result in a 2D scatter plot. The individual “spot” fluorescence data is given in Supp. Table 1.

### Transposon-wrmScarlet reporter

This reporter was constructed by fusing the wrmScarlet at the 3’end, just before the translation stop codon of ZC15.3, that contains a pao class LTR retrotransposon element. Required DNA fragments were generated as following: PCR with RG1134-RG932 (template is gDNA), RG1135-RG1136 (amplifying the wrmScarlet), RG1137-RG912 (template is gDNA). A gibson reaction was performed by mixing these three PCR products and AvrII/BglII linearized pCFJ150 plasmid generating wrmScarlet tagged ZC15.3.

### RAD-51 staining

L4 staged animals were used and staining was done essentially as described in www.wormatlas.org/images/finneyruvkun.pdf. with the below mentioned modifications. Once the worms were fixed, the worms were opened by Freeze-cracking. Rabbit anti-RAD-51 Antibodies (Novu Biologics, 2948002) at a dilution 1:500 and secondary antibodies (Goat anti-Rabbit IgG (H+L) Highly Cross-Adsorbed Secondary Antibody, Alexa Fluor 488, Invitrogen, A-11034) at a concentration of 1:1000 was used. Confocal imaging was done as described earlier. The animals that showed RAD-51 foci in both mitotic & transition zone and “late pachytene zone” are included in the analysis.

### Preparing the worm extract

Worms grown at indicated temperature and of defined developmental stages were collected and washed three times in M9 buffer. The worms were transferred to 2 ml tubes (Sarstedt, Ref: 72.693.005) with Zirconia/Silica beads (Biospec, Cat. No. 110791052) in lysis buffer (50mM Tris-HCl, pH 7.5, 150 mM NaCl, 1% TRITON X-100, 1 mM EDTA) supplemented with 1mM PMSF and 1 tablet of cOmplete Protease inhibitor (Roche, cat. No. 11 873 580 001) per 50 ml. The worms were lysed using MP Biomedicals Fast Prep-24 5G bead beater for 16 cycles at 8m/Sec. The lysate was centrifuged at 16100g for 10 minutes at 4°C and the supernatant was transferred to a fresh tube and the concentration was measured using standard bradford assay.

### Immunoprecipitation of tagged proteins of interest

For Immunoprecipitation (IP), the cleared worm extract was incubated with 50ul of 50% bead suspension of anti-flag M2 Magnetic Beads (Sigma. Catalog Number M8823) that were pre-washed thrice with TBS buffer (50 mM Tris-HCl pH 7.5, 150 mM NaCl) on a rotating wheel at 4°C for 2 Hours. The beads were then washed once with lysis buffer that contained protease inhibitors and then washed 6 more times with TBS buffer without protease inhibitors.

### On beads digestion

After the IP is done, 5ul of digestion buffer (3M GuaHCl, 20mM EPPS pH8.5, 10mM CAA, 5mM TCEP) and 1ul of Lys-C (0.2ug/ul in 50mM Hepes, pH8.5) is added and incubated at 21°C for 2-4 Hours while shaking. 17 ul of 50mM Hepes, pH8.5 is added. 1ul of 0.2ug/ul trypsin is added and incubated at 37°C overnight. In the morning, another ul of trypsin is added and incubated for 4 more hours and Mass Spectrometry is done as described below.

### Small RNA sequencing following Immunoprecipitation

IP was done exactly as described above with the exception that RNAse inhibitor (Promega, N261A) at 200U/IP was used during the first 2 Hrs of the IP. Following IP, the RNA extraction and library preparation was done as described earlier.

### Mapping the *in vivo* cleavage sites of WAGO-1 and WAGO-3

Once the protein of interest is IPed from the desired genetic background, the protein was “on beads” digested as described earlier. We have used L4 staged worms for this experiment. For WAGO-1, the “on beads” digestion protocol was followed exactly as described. For WAGO-3, in addition to trypsin, AspN enzyme was used.

### *in vitro* protease assay and Mass Spectrometry

The wild type DPF-3 or *DPF-3(S784A)* mutant protein was IPed from staged worm extract as described earlier. The “Beads only” control without worm extract is prepared simultaneously. The peptides were ordered at Thermo Fischer Scientific and the stock solution of peptides was made by dissolving the lyophilized peptides in TBS buffer supplemented with 2mm TCEP to get a final concentration of 0.1 nmol/ul. we pooled both the peptides at a final concentration of 1pmol/ul of each peptide. The “beads only” control or the beads with bound protein of interest is then incubated with pooled peptides at 20°C, aliquots were taken out at the indicated time and mixed with 180 ul of 0.1% trifluoroacetic acid, 2mM TCEP, 2% acetonitrile in water to stop the reaction. The peptides were then analyzed by capillary liquid chromatography tandem mass spectrometry (LC-MS). Peptides were loaded onto a 75 uml5 cm ES811 column (Accucore C4, 2.6□μm, 150□Å) at a constant pressure of 800 bar, using an EASY-nLC 1000 liquid chromatograph with one-column set up (Thermo Scientific). Peptides were separated at a flow rate of 300□nl/min with a linear gradient of 2–6% buffer B in buffer A in 2□min, followed by a linear increase from 6-22% in 20□min, 22%-28% in 4 min, 28-36% in 2min, 36-80% in 1min and the column was finally washed for 5□min at 90% buffer B (buffer A: 0.1% formic acid in water; buffer B: 0.1% formic acid in acetonitrile). The column was mounted on a DPV ion source (New Objective) connected to an Orbitrap Fusion mass spectrometer (Thermo Scientific), data were acquired using 120,000 resolution, for the peptide measurements in the Orbitrap and a top T (1.5□s) method with HCD fragmentation for each precursor and fragment measurement in the ion trap following the manufacturer guidelines (Thermo Scientific). MS1 signals were quantified using Skyline 4.1 (MacLean *et al*., 2010) to generate the results Fig. 6A.

Tryptic peptides of Immunoprecipitated WAGO-1 were analyzed by LC–MS/MS, essentially as described (Ostapcuk *et al*., 2018). In short, the peptides were separated with an EASY-nLC 1000 on a 50 μm□×□15□cm ES801 C18, 2 μm, 100□Å column (Thermo Scientific) mounted on a DPV ion source (New Objective). They were measured with a Orbitrap Fusion (Thermo Scientific) using a top T (3□s) method as recommended by the manufacturer (Thermo Scientific). Andromeda implemented in MaxQuant (version: 1.5.3.8) (Cox *et al*., 2011) was used to search the CAEEL subset of the UniProt (version: 2017_04) combined with the contaminant database from MaxQuant and label-free quantification (LFQ) (Cox *et al*., 2014) was used with a protein and peptide FDR of 0.01. Statistical analysis was done in Perseus (version: 1.5.2.6) (Hubner *et al*., 2010; Tyanova *et al*., 2016). MS1 signals of selected WAGO-1 and WAGO-3 peptides were quantified using Skyline 4.1 (MacLean *et al*., 2010) to generate the results in Fig. 6B and fig. S6D, E.

### Peptide description

For peptides mapping to WAGO-1 (Uniprot ID Q21770), these were generated by digesting with trypsin that cleaves C-terminus of lysine or arginine. “S2” peptide represents the sequence present between S2-K37, “H4” represents H4-K37 etc. The internal peptides (V267: V267-R380; S516: S516-K523; A820: A820-K841) are coming from region farther to N-terminus. Regarding the peptides mapping to WAGO-3, these were generated by digesting with trypsin and AspN enzymes. The first cleavage in WAGO-3 ((Uniprot ID Q9N585), would happen just before the FLAG tag (DYKDDDDK) inserted between S46 and E47, hence the “P2” peptide represents sequence between P2 and S46, “V6” represents V6-S46 etc. The internal peptides (D237: D237-K248; D571: D571-R578; G709: G709-K725) are coming from region farther to N-terminus.

### Antigen for anti DPF-3 antibodies

The C-terminal region of DPF-3 (amino acids 641-931) was used as antigen to raise the antibodies against *C. elegans* DPF-3 in rats. The antigen amino acid sequence is given below.

TVYAANITVSGHPGQPDLHFDSPEMIEFQSKKTGLMHYAMILRPSNFDPYKKYPVFHYVYGGPGIQIVHNDFSWIQYIRFCRLGYVVVFIDNRGSAHRGIEFERHIHKKMGTVEVEDQVEGLQMLAERTGGFMDMSRVVVHGWSYGGYMALQMIAKHPNIYRAAIAGGAVSDWRLYDTAYTERYMGYPLEEHVYGASSITGLVEKLPDEPNRLMLVHGLMDENVHFAHLTHLVDECIKKGKWHELVIFPNERHGVRNNDASIYLDARMMYFAQQAIQGFGPTTAAPRQGPL

### Western blot

Total worm extract was loaded in equal quantities. Primary antibody against DPF-3 was used at 1:200 dilution and anti-rat secondary antibodies fused to HRP (Jackson ImmunoResearch, 712-035-150) were used at 1:2000 dilution. Amersham ECL Prime western blotting reagents were used and the bands were detected with an ImageQuant LAS 4000 chemiluminescence imager.

### Pharmacological inhibition

The animals containing the fluorescent reporter (HW2283: xeSi410[Pzc15.3::ZC15.3-Wrm Scarlet::ZC15.3 3’UTR,, unc-119(+)]III) were grown on NG2% plates supplemented with OP50 bacteria. The embryos were extracted from gravid adults by bleaching and the eggs were introduced into the well of a corning plate (CLS3513, Sigma Aldrich) containing OP50 at an OD of 1.0 diluted in S-basal medium (For 1 Liter, 5.85 g of NaCl, 1 g of K_2_HPO_4_, 6 g of KH_2_PO_4_, 1 ml of 5mg/ml Cholesterol dissolved in ethanol). 1G244 (SML2247) and Vildagliptin (SML2302) purchased from Sigma Aldrich were dissolved in DMSO to make stock solutions at 10mM and 100mM respectively. Pure DMSO was used as a control. Indicated concentration of drug was added to the wells and incubated at 26.5°C and the animals were observed for the de-silencing of the fluorescence reporter or were collected 42 Hrs later for total RNA extraction as described earlier.

### Phylogenetic tree

The phylogenetic tree of the DPF proteins was constructed using http://www.phylogeny.fr/index.cgi as described in (Dereeper *et al*., 2008).

### High Throughput Sequencing data analysis

Processing of the High Throughput Sequencing libraries was done using snakemake (Koster and Rahmann, 2018) pipelines. Conda (https://conda.io/) was used to install the necessary dependencies mainly via the bioconda (Gruning *et al*., 2018) channel. The snakemake pipelines, custom analyses scripts and the jupyter notebooks (Kluyver *et al*., 2016) used to generate the final plots are available in https://github.com/fmi-basel/ggrosshans_analysis_dpf-3. (*Currently private but for review accessible here*: *https://www.dropbox.com/s/bklvdet6mvn7xjz/ggrosshans_analysis_dpf-3-master.zip?dl=0*) The specific versions of the tools used are also available in the repository.

### Annotation

The *C. elegans* genome and the corresponding annotation files were obtained from WormBase (Harris *et al*., 2020) (release WS270, ftp://ftp.wormbase.org/pub/wormbase/releases/WS270/species/c_elegans/PRJNA13758/). Genes from the canonical gene set (c_elegans.PRJNA13758.WS270.canonical_geneset.gtf) were used. Repeats (transposons are also included in this annotation) were obtained from the table browser of UCSC (Karolchik *et al*., 2004) after selecting as group “Variation and Repeats” and as track “RepeatMasker”.

In order to use both genes and repeats in the different analyses, we constructed a hybrid gene/repeat annotation. Repeats of Class Simple_repeat, Low_complexity or rRNA were excluded. With the aid of bedtools (Quinlan and Hall, 2010) and custom scripts that rely on pandas (McKinney, 2011) and HTSeq (Anders, Pyl and Huber, 2015) we intersected the exons of the genes with the filtered repeats. Exons that overlapped with repeats (on the same strand) were excluded. The final transcriptome contains transcripts that do not overlap with repeats (in exonic regions of the same strand), transcripts that overlap with repeats (overlapping exons were excluded) and repeats. Genome indexes for bowtie (Langmead *et al*., 2009) and STAR (Dobin *et al*., 2013) were generated. The pipeline to generate the custom annotation and the indexes is available in the directory 00_annotation of the repository.

### Analysis of small RNA-seq

Quality of the reads were examined with FastQC (Andrews and Others, 2010). With the aid of custom scripts that rely on HTSeq (Anders, Pyl and Huber, 2015), pandas (McKinney, 2011) and BioPython (Cock *et al*., 2009) we keep the reads that contain at least 8 bases after the 3’adapter (AACTGTAGGCACCATCAAT), because the UMIs are incorporated after the 3’adapter. The 3’ adapters were removed with cutadapt (Martin, 2011) (--error-rate 0.1, --minimum-length 15, --overlap 3), low quality reads were filtered with fastq_quality_filter from FASTX-Toolkit (http://hannonlab.cshl.edu/fastx_toolkit/) (-q 20, -p 100, -Q 33) and reads are then collapsed based on the UMIs. Reads were mapped to the genome with bowtie (Langmead *et al*., 2009) (-v 0, --all, --best, --strata) and converted to bam files with samtools (Li *et al*., 2009). A custom script was used to filter reads that fall in transcripts annotated (transcript_biotype) as rRNA, rRNA_pseudogene, tRNA, tRNA_pseudogene. Reads from MtDNA were also filtered. Filtered reads were converted back to fastq files with a custom script. Reads were mapped again to the genome with bowtie (Langmead *et al*., 2009) (-v 0, --all, -M 1, --best, --strata). Genome coverage tracks were generated with bedtools (Quinlan and Hall, 2010) and converted to bigwig files with bedGraphToBigWig (Kent *et al*., 2010). ALFA (Bahin *et al*., 2019) was used to determine the different biotypes that the reads map to. We used htseq-count from HTSeq (Anders, Pyl and Huber, 2015) to count reads that fall in the opposite strand of reads/repeats (--stranded reverse, --type exon, --idattr gene_id, --mode union, -a 0, --nonunique none, --secondary-alignments ignore, --supplementary-alignments ignore). Only transcripts annotated as protein_coding, or repeat were quantified. Differential expression analysis was performed with edgeR (Robinson, McCarthy and Smyth, 2010) with filterByExpr, glmFit, glmLRT and p.adjust(…,method=“BH”) (Benjamini and Hochberg, 1995). Reads were also counted at the sequence level with a custom script. This means that instead of counting the reads that fall in the opposite strand of genes or repeats we used the read sequence as the identifier. This allowed us to separate different small RNA categories (22G RNAs, 21U RNAs). Sequences of mature miRNAs were excluded from this analysis. The pipeline is available in the directory 01_small_RNA_seq_15_C of the repository.

### Analysis of total RNA-seq

3’adapters (AGATCGGAAGAGCACACGTCTGAACTCCAGTCA, AGATCGGAAGAGCGTCGTGTAGGGAAAGAGTGT) were trimmed with cutadapt (Martin, 2011) and similar parameters as above. Reads were mapped to the genome with STAR (Dobin *et al*., 2013) (default parameters), alignment files were converted to wiggle files with STAR (Dobin *et al*., 2013) and bigwig files were generated with wigToBigWig (Kent *et al*., 2010). Reads were counted with featureCounts (Liao, Smyth and Shi, 2014) (-M -O --fraction -s 2 -p -B -P -C). Differential expression analysis was performed with edgeR (Robinson, McCarthy and Smyth, 2010) (as previously). The pipeline is available in the directory 02_total_RNAseq_15_C of the repository.

### Analysis of IPs

Reads were processed in a similar manner as the small RNA-seq. The main difference is that 12 bases after the 3’adapter (AACTGTAGGCACCATCAAT) were used as UMI. A snakemake pipeline is available in the directory 03_WAGO_IPs and 04_csr1_ip_and_wago_double_mutants of the repository.(Ishidate *et al*., 2018)

### Plotting

The scripts that generate the different plots are under the directory plots of the repository. These rely on the libraries pandas(McKinney, 2011), numpy (Van Der Walt, Colbert and others, 2011), seaborn (Waskom *et al*., 2014) and gviz (Hahne and Ivanek, 2016).

---

**Fig. S1.**
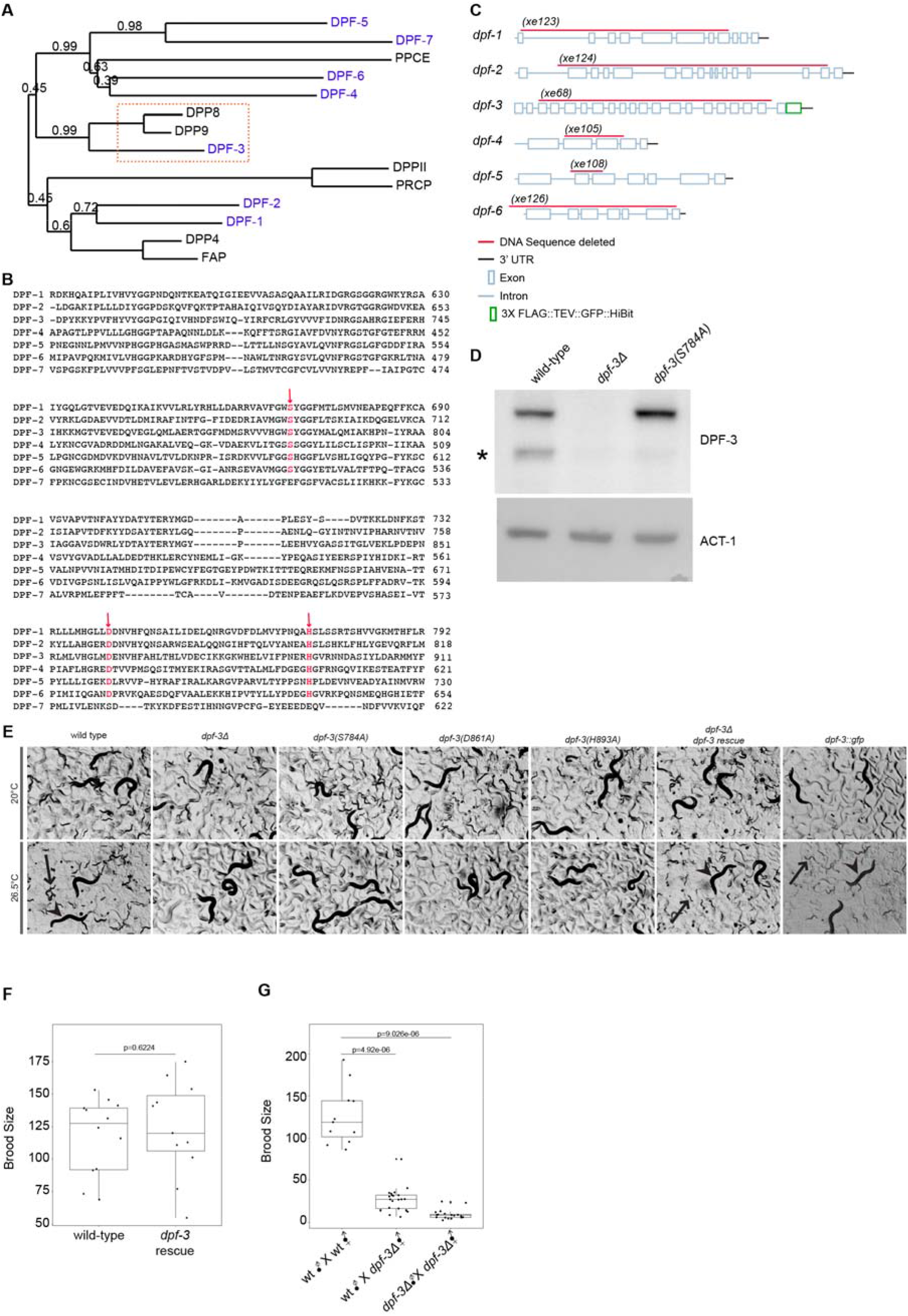
*dpf-3* is required for fertility. **A.** Phylogenetic tree of DPPIV family proteins of *H. sapiens* (black) and *C. elegans* (blue). A red dotted rectangle highlights the orthologous group of *C. elegans* DPF-3 and human DPP8/DPP9. Numbers indicate bootstrap values. **B**. Sequence alignment of the peptidase domain of the DPPIV family proteins generated by Clustal Omega (Madeira *et al*., 2019). The three amino acids constituting the catalytic triad are colored red and indicated with a red arrow pointing down. Note the conservation of the residues in all DPFs except DPF-7. **C**. Schematic depiction of *dpf* gene architectures, approximately to scale. Red lines indicate deletions used in the study, a green box the GFP tag inserted just before the stop codon of *dpf-3*. **D**. Western blot probed with anti-DPF-3 antibodies showing the accumulation of DPF-3 in wild-type and *dpf-3(S784A)* L1 stage animals (grown at 25°C for 10 h after plating synchronized larvae on food). ACT-1 serves as a loading control. An asterisk indicates an N-terminally truncated form of DPF-3 that is regularly observed, but which we did not investigate further. **E**. Representative micrographs of animals grown at the indicated temperatures reveal temperature-dependent fertility defects of *dpf-3* mutant animals. Where present, individual P0 and F1 progeny are indicated with arrowheads and arrows respectively. No viable larvae were observed in the *dpf-3* mutant animals at 26.5°C. *dpf-3::gfp* is the *dpf-3(xe246)* (see below) strain with endogenously tagged *dpf-3*. **F.** Box plot of the brood size of wild-type and *dpf-3* rescue animals grown at 26. 5°C. **G.** Box plot of the brood size of two different crosses. The males and hermaphrodites were grown at 20°C and 26.5°C respectively. Crossing was performed at 26.5°C. p values in **F** and **G** are calculated using a Wilcoxon test.

**fig. S2.**
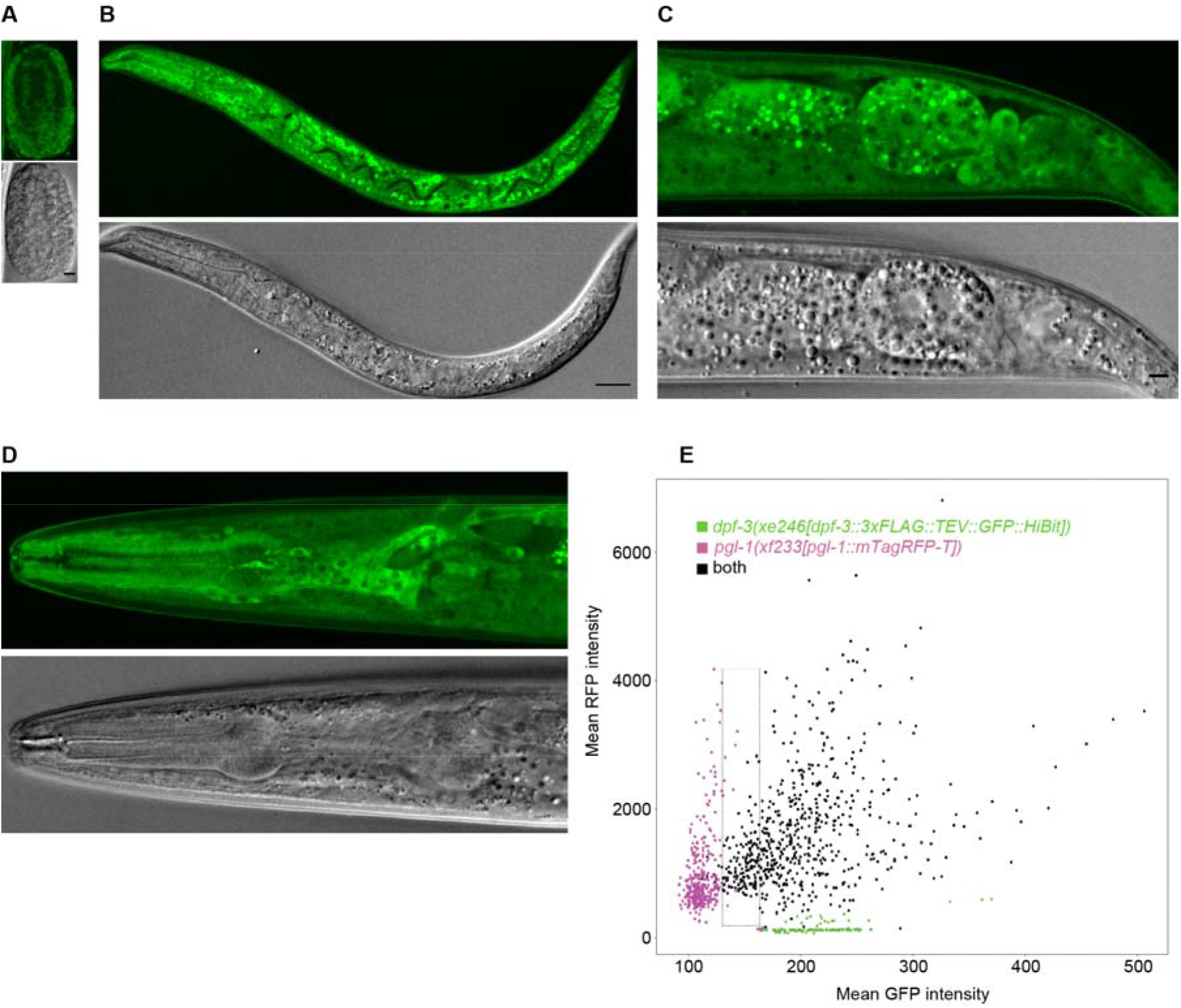
*dpf-3* is broadly expressed. **A-D.** Single plane confocal images showing *dpf-3* expression in embryo (A), whole L1 larva (B), adult tail (C) and head (D). Scale bars are 10 μm. **E.** Scatter plot showing the co-localization of DPF-3 (green) and PGL-1 (purple) observed in Fig. 2C. Each dot represents a single punctum observed in developing oocytes located between the gonad bend and the uterus of staged young adult animals. The x- and the y-axis indicate the mean GFP and RFP intensity respectively. The green and purple dots are from strains expressing only *dpf-3::gfp* and only *pgl-1::rfp*, respectively, revealing sensitivity and specificity of signal detection from each fluorophore. The black dots are from a strain expressing both tagged genes. Although most of the DPF-3::GFP punctae are also occupied by PGL-1::RFP, the converse is not true, and a dotted rectangle encompasses punctae with PGL-1 but not DPF-3 signal above background. The fluorescence intensity of each punctum or “spot” is given in the Supp. table 1

**Fig. S3.**
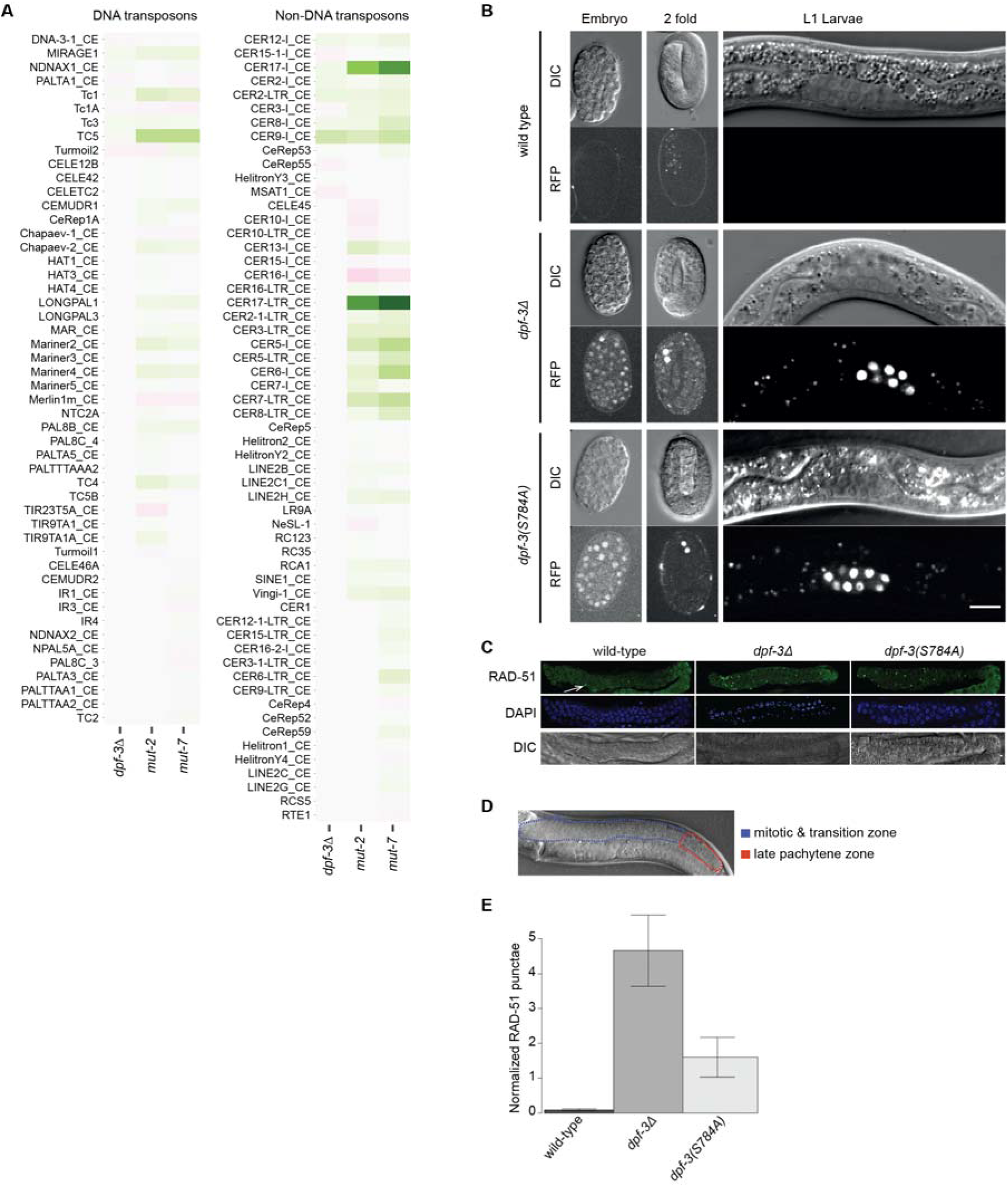
Loss of *dpf-3* leads to transposon de-silencing. **A**. Heatmap showing different DNA and non-DNA transposons (y-axis) that are significantly changed in the indicated mutant relative to wild-type animals (x-axis). **B.** Single plane DIC and fluorescent confocal images of different developmental stages of wild-type and *dpf-3* mutant animals expressing the *ZC15.3*::wrmScarlet transgene grown at 15°C. Scale bar is 10 μm. **C.** Representative images of fixed gonads from late L4 stage animals of the indicated genotypes grown at 25°C. The white arrow points to a RAD-51 punctum, visualized by anti-RAD-51 immunofluorescent staining. **D.** Contours showing the approximate mitotic + transition zone (blue) and late pachytene zone (red) of a late L4 stage animal. **E.** Quantification of RAD-51 punctae in mitotic + transition zone in animals of the indicated genotypes. To control for staining efficiency, the number is normalized to number of punctae in late pachytene zone, where meiotic recombination causes a natural occurrence of RAD-51 punctae. The average of wild-type (3 animals) and *dpf-3* mutants (6 animals of each mutant) is shown. Error bars indicate Standard Error of Mean (SEM).

**Fig. S4.**
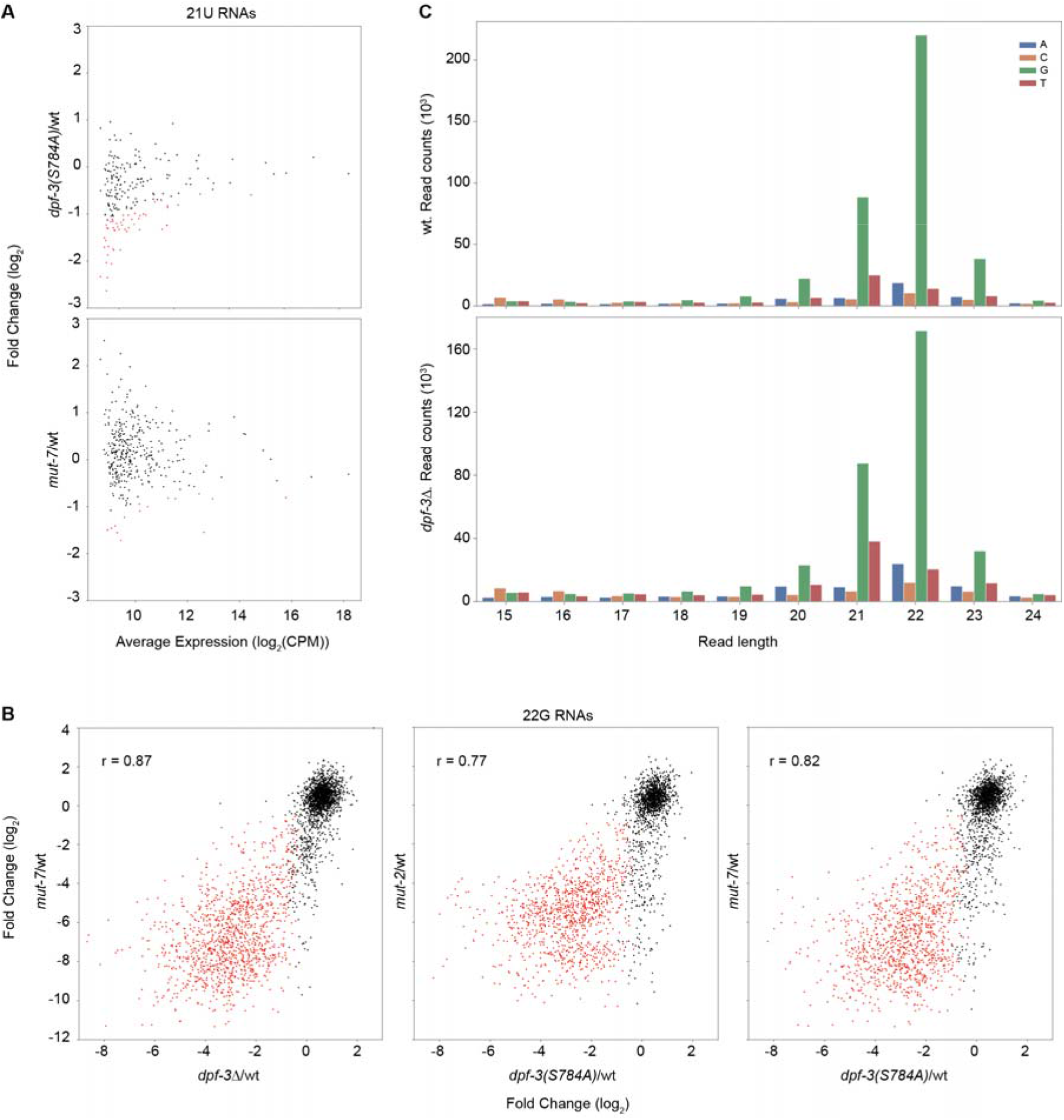
Specific 22G RNAs are down-regulated in *dpf-3* mutant animals. **A, B.** Scatter plots showing differential expression of 21U RNAs (A) or 22G RNAs (B) between wild-type and the indicated mutant animals. Each dot represents an individual 21U RNA (A) or all the 22G RNAs that match in anti-sense orientation to a given transcript (B); red indicates a significant change (FDR<0.05). The r values indicate the Pearson correlation coefficient. **C.** Read length distribution of small RNAs starting with a given base indicated with different colors for wild-type (top) and *dpf-3Δ* mutant (bottom) samples.

**Fig. S5.**
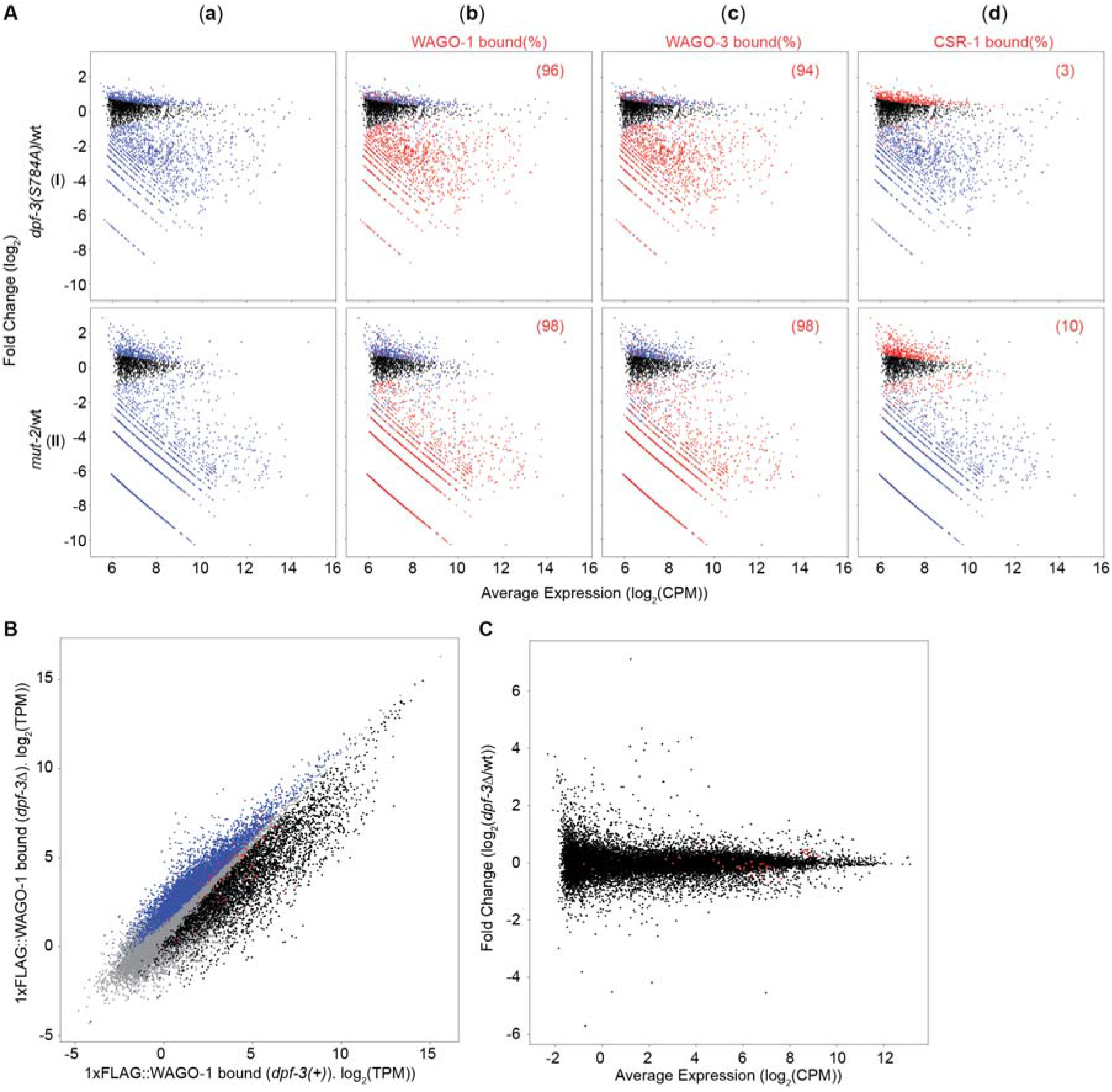
Most of the 22G RNAs lost in dpf-3 mutant animals are bound by WAGO-1 and WAGO-3. **A**. MA plots showing the differential expression of 22G RNAs in the indicated mutant relative to wild-type animals (rows I-II) plotted over average expression in the respective mutant and wild-type condition (as CPM – Counts Per Million mapped reads). Each dot represents all the 22G RNAs that match in anti-sense orientation to a given transcript. (**a**) Significantly changing (FDR<0.05) transcripts are colored blue. (**b**-**d**) The 22G RNAs that are bound by a specific Argonaute protein (and differentially expressed in *dpf-3* and *mut-2* mutant animals are colored red. The number (%) shown in each individual plot indicates the fraction of 22G RNAs bound by a specific Argonaute protein and down regulated in indicated mutant animals. **B**. Scatter plot from Fig. 5B showing binding of 22G RNAs to WAGO-1 in lysates from *dpf-3Δ* (y-axis) and wild-type (x-axis) L4 stage animals, with histone genes colored in red. Gray indicates no change, black significant change depletion and blue significant enrichment on WAGO-1 *in dpf-3* mutant relative to wild-type animals (FDR<0.05). **C**. MA plot showing differential expression (in log_2_) of transcripts (total RNA) in the *dpf-3Δ* mutant relative to wild-type animals plotted over average expression in the respective mutant and wild-type condition (as CPM – Counts Per Million mapped reads). Each dot represents an individual transcript; transcripts coding for histone proteins are colored red.

**Fig. S6.**
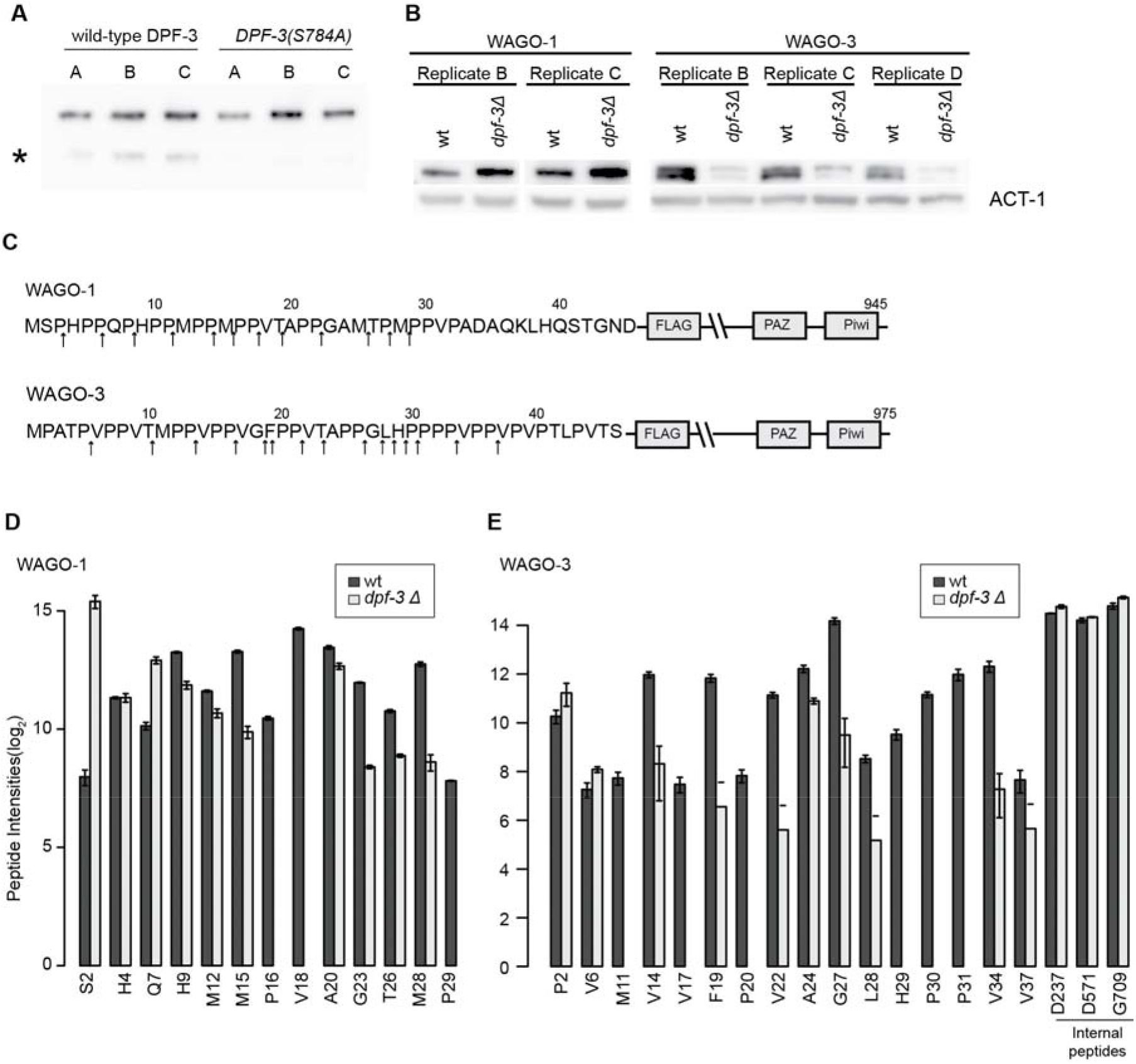
WAGO-1 and WAGO-3 are processed by DPF-3. **A.** Western blot probed with anti-FLAG antibodies revealing similar levels of wild type and S784A mutated *DPF-3* in immunoprecipitated samples used for the *in vitro* protease assay shown in (Fig. 6A). The biological triplicates are denoted as A, B, C. The lower band indicated with * represent the N-terminally truncated form of DPF-3. **B**. Biological replicates of western blot described in Fig. 6C. **C.** Schematic depiction of WAGO-1 and WAGO-3 proteins showing the transgenic FLAG tag and the PAZ & Piwi domains (not to scale). The number on the top indicates the amino acid position in the cognate protein. The arrows pointing up indicate the *in vivo* cleavage positions observed. **D-E.** Bar plot showing the log_2_-transformed average peptide intensities of peptides starting in the indicated amino acid derived from WAGO-1 (D) and WAGO-3 (E) for three biological replicates. For details about the different peptides labelled on the x-axis, see “peptide description” in materials and methods section. The “S2” peptide data is replotted from Fig. 6B for reference. Error bars represent Standard Error of Mean (SEM).

**fig. S7.**
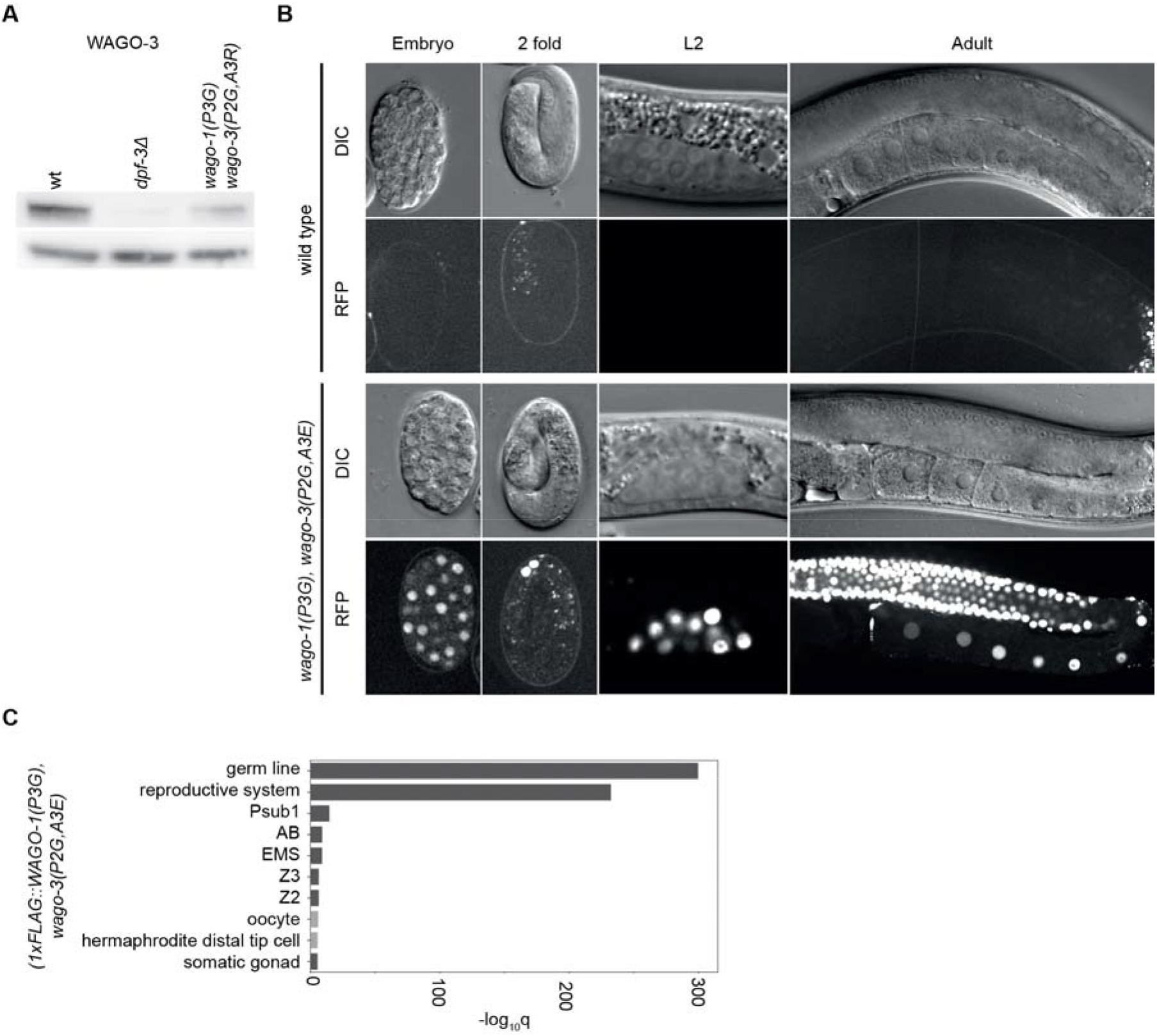
Mutation of *wago-1* and *wago-3* leads to de-silencing of transposons. **A**. Biological replicate of the western blot shown in Fig. 7B. **B**. Single plane DIC and fluorescent confocal images of different developmental stages of wild-type and *wago-1(P3G), wago-3(P2G,A3E)* mutant animals expressing the *ZC15.3*::wrmScarlet transgene grown at 15°C. The images of wild-type are from Fig. 3D and fig. S3B and included for comparison. **C**. Tissue Enrichment Analysis (TEA) of 22G RNAs bound to WAGO-1 in *wago-1(P3G), wago-3(P2G,A3E)* mutant animals (blue points from (Fig. 7D)). Tissues that are commonly enriched in *wago-1(P3G), wago-3(P2G,A3E)* and *dpf-3Δ* mutant animals (Fig. 5D) are colored dark grey. The x-axis indicates *q* values (-log_10_).

## Notes

### Competing Interest Statement

The authors have declared no competing interest.

## References

Aeschimann, F., Kumari, P., Bartake, H., Gaidatzis, D., Xu, L., Ciosk, R. and Grosshans, H. (2017) ‘LIN41 Post-transcriptionally Silences mRNAs by Two Distinct and Position-Dependent Mechanisms’, Mol Cell, 65(3), pp. 476–489 e4.

Ajami, K., Abbott, C. A., Obradovic, M., Gysbers, V., Kahne, T., McCaughan, G. W. and Gorrell, M. D. (2003) ‘Structural requirements for catalysis, expression, and dimerization in the CD26/DPIV gene family’, Biochemistry, 42(3), pp. 694–701.

Almeida, M. V., de Jesus Domingues, A. M. and Ketting, R. F. (2019) ‘Maternal and zygotic gene regulatory effects of endogenous RNAi pathways’, PLoS Genet, 15(2), pp. e1007784.

Anders, S., Pyl, P. T. and Huber, W. (2015) ‘HTSeq--a Python framework to work with high-throughput sequencing data’, Bioinformatics, 31(2), pp. 166–169.

Andrews, S. and Others (2010) ‘FastQC: a quality control tool for high throughput sequence data’.

Arribere, J. A., Bell, R. T., Fu, B. X., Artiles, K. L., Hartman, P. S. and Fire, A. Z. (2014) ‘Efficient marker-free recovery of custom genetic modifications with CRISPR/Cas9 in Caenorhabditis elegans’, Genetics, 198(3), pp. 837–46.

Bahin, M., Noel, B. F., Murigneux, V., Bernard, C., Bastianelli, L., Le Hir, H., Lebreton, A. and Genovesio, A. (2019) ‘ALFA: annotation landscape for aligned reads’, BMC Genomics, 20(1), pp. 250.

Barucci, G., Cornes, E., Singh, M., Li, B., Ugolini, M., Samolygo, A., Didier, C., Dingli, F., Loew, D., Quarato, P. and Cecere, G. (2020) ‘Small-RNA-mediated transgenerational silencing of histone genes impairs fertility in piRNA mutants’, Nat Cell Biol, 22(2), pp. 235–245.

Batista, P. J., Ruby, J. G., Claycomb, J. M., Chiang, R., Fahlgren, N., Kasschau, K. D., Chaves, D. A., Gu, W., Vasale, J. J., Duan, S., Conte, D., Jr., Luo, S., Schroth, G. P., Carrington, J. C., Bartel, D. P. and Mello, C. C. (2008) ‘PRG-1 and 21U-RNAs interact to form the piRNA complex required for fertility in C. elegans’, Mol Cell, 31(1), pp. 67–78.

Benjamini, Y. and Hochberg, Y. (1995) ‘Controlling the False Discovery Rate: A Practical and Powerful Approach to Multiple Testing’, Journal of the Royal Statistical Society: Series B (Methodological), 57(1), pp. 289–300.

Brown, K. C., Svendsen, J. M., Tucci, R. M., Montgomery, B. E. and Montgomery, T. A. (2017) ‘ALG-5 is a miRNA-associated Argonaute required for proper developmental timing in the Caenorhabditis elegans germline’, Nucleic Acids Res, 45(15), pp. 9093–9107.

Burkey, B. F., Hoffmann, P. K., Hassiepen, U., Trappe, J., Juedes, M. and Foley, J. E. (2008) ‘Adverse effects of dipeptidyl peptidases 8 and 9 inhibition in rodents revisited’, Diabetes Obes Metab, 10(11), pp. 1057–61.

Chen, C. C., Simard, M. J., Tabara, H., Brownell, D. R., McCollough, J. A. and Mello, C. C. (2005) ‘A member of the polymerase beta nucleotidyltransferase superfamily is required for RNA interference in C. elegans’, Curr Biol, 15(4), pp. 378–83.

Claycomb, J. M., Batista, P. J., Pang, K. M., Gu, W., Vasale, J. J., van Wolfswinkel, J. C., Chaves, D. A., Shirayama, M., Mitani, S., Ketting, R. F., Conte, D., Jr. and Mello, C. C. (2009) ‘The Argonaute CSR-1 and its 22G-RNA cofactors are required for holocentric chromosome segregation’, Cell, 139(1), pp. 123–34.

Cock, P. J., Antao, T., Chang, J. T., Chapman, B. A., Cox, C. J., Dalke, A., Friedberg, I., Hamelryck, T., Kauff, F., Wilczynski, B. and de Hoon, M. J. (2009) ‘Biopython: freely available Python tools for computational molecular biology and bioinformatics’, Bioinformatics, 25(11), pp. 1422–3.

Conine, C. C., Batista, P. J., Gu, W., Claycomb, J. M., Chaves, D. A., Shirayama, M. and Mello, C. C. (2010) ‘Argonautes ALG-3 and ALG-4 are required for spermatogenesis-specific 26G-RNAs and thermotolerant sperm in Caenorhabditis elegans’, Proc Natl Acad Sci U S A, 107(8), pp. 3588–93.

Conine, C. C., Moresco, J. J., Gu, W., Shirayama, M., Conte, D., Jr., Yates, J. R., 3rd and Mello, C. C. (2013) ‘Argonautes promote male fertility and provide a paternal memory of germline gene expression in C. elegans’, Cell, 155(7), pp. 1532–44.

Cox, J., Hein, M. Y., Luber, C. A., Paron, I., Nagaraj, N. and Mann, M. (2014) ‘Accurate proteome-wide label-free quantification by delayed normalization and maximal peptide ratio extraction, termed MaxLFQ’, Mol Cell Proteomics, 13(9), pp. 2513–26.

Cox, J., Neuhauser, N., Michalski, A., Scheltema, R. A., Olsen, J. V. and Mann, M. (2011) ‘Andromeda: a peptide search engine integrated into the MaxQuant environment’, J Proteome Res, 10(4), pp. 1794–805.

de Albuquerque, B. F., Placentino, M. and Ketting, R. F. (2015) ‘Maternal piRNAs Are Essential for Germline Development following De Novo Establishment of Endo-siRNAs in Caenorhabditis elegans’, Dev Cell, 34(4), pp. 448–56.

Deacon, C. F. (2019) ‘Physiology and Pharmacology of DPP-4 in Glucose Homeostasis and the Treatment of Type 2 Diabetes’, Front Endocrinol (Lausanne), 10, pp. 80.

Delaney, K., Mailler, J., Wenda, J. M., Gabus, C. and Steiner, F. A. (2018) ‘Differential Expression of Histone H3.3 Genes and Their Role in Modulating Temperature Stress Response in Caenorhabditis elegans’, Genetics, 209(2), pp. 551–565.

Dereeper, A., Guignon, V., Blanc, G., Audic, S., Buffet, S., Chevenet, F., Dufayard, J. F., Guindon, S., Lefort, V., Lescot, M., Claverie, J. M. and Gascuel, O. (2008) ‘Phylogeny.fr: robust phylogenetic analysis for the non-specialist’, Nucleic Acids Research, 36, pp. W465–W469.

Dobin, A., Davis, C. A., Schlesinger, F., Drenkow, J., Zaleski, C., Jha, S., Batut, P., Chaisson, M. and Gingeras, T. R. (2013) ‘STAR: ultrafast universal RNA-seq aligner’, Bioinformatics, 29(1), pp. 15–21.

Dokshin, G. A., Ghanta, K. S., Piscopo, K. M. and Mello, C. C. (2018) ‘Robust Genome Editing with Short Single-Stranded and Long, Partially Single-Stranded DNA Donors in Caenorhabditis elegans’, Genetics, 210(3), pp. 781–787.

Frokjaer-Jensen, C., Davis, M. W., Ailion, M. and Jorgensen, E. M. (2012) ‘Improved Mos1-mediated transgenesis in C. elegans’, Nat Methods, 9(2), pp. 117–8.

Gent, J. L., Lamm, A. T., Pavelec, D. M., Maniar, J. M., Parameswaran, P., Tao, L., Kennedy, S. and Fire, A. Z. (2010) ‘Distinct phases of siRNA synthesis in an endogenous RNAi pathway in C. elegans soma’, Mol Cell, 37(5), pp. 679–89.

Gibson, D. G., Young, L., Chuang, R. Y., Venter, J. C., Hutchison, C. A. and Smith, H. O. (2009) ‘Enzymatic assembly of DNA molecules up to several hundred kilobases’, Nature Methods, 6(5), pp. 343–U41.

Gruning, B., Dale, R., Sjodin, A., Chapman, B. A., Rowe, J., Tomkins-Tinch, C. H., Valieris, R., Koster, J. and Bioconda, T. (2018) ‘Bioconda: sustainable and comprehensive software distribution for the life sciences’, Nat Methods, 15(7), pp. 475–476.

Gu, W., Shirayama, M., Conte, D., Jr., Vasale, J., Batista, P. J., Claycomb, J. M., Moresco, J. J., Youngman, E. M., Keys, J., Stoltz, M. J., Chen, C. C., Chaves, D. A., Duan, S., Kasschau, K. D., Fahlgren, N., Yates, J. R., 3rd, Mitani, S., Carrington, J. C. and Mello, C. C. (2009) ‘Distinct argonaute-mediated 22G-RNA pathways direct genome surveillance in the C. elegans germline’, Mol Cell, 36(2), pp. 231–44.

Hahne, F. and Ivanek, R. (2016) ‘Visualizing Genomic Data Using Gviz and Bioconductor’, Methods Mol Biol, 1418, pp. 335–51.

Harris, T. W., Arnaboldi, V., Cain, S., Chan, J., Chen, W. J., Cho, J., Davis, P., Gao, S., Grove, C. A., Kishore, R., Lee, R. Y. N., Muller, H. M., Nakamura, C., Nuin, P., Paulini, M., Raciti, D., Rodgers, F. H., Russell, M., Schindelman, G., Auken, K. V., Wang, Q., Williams, G., Wright, A. J., Yook, K., Howe, K. L., Schedl, T., Stein, L. and Sternberg, P. W. (2020) ‘WormBase: a modern Model Organism Information Resource’, Nucleic Acids Res, 48(D1), pp. D762–D767.

Hauptmann, J., Dueck, A., Harlander, S., Pfaff, J., Merkl, R. and Meister, G. (2013) ‘Turning catalytically inactive human Argonaute proteins into active slicer enzymes’, Nat Struct Mol Biol, 20(7), pp. 814–7.

Hubner, N. C., Bird, A. W., Cox, J., Splettstoesser, B., Bandilla, P., Poser, L., Hyman, A. and Mann, M. (2010) ‘Quantitative proteomics combined with BAC TransgeneOmics reveals in vivo protein interactions’, J Cell Biol, 189(4), pp. 739–54.

Ishidate, T., Ozturk, A. R., Durning, D. J., Sharma, R., Shen, E. Z., Chen, H., Seth, M., Shirayama, M. and Mello, C. C. (2018) ‘ZNFX-1 Functions within Perinuclear Nuage to Balance Epigenetic Signals’, Mol Cell, 70(4), pp. 639–649 e6.

Justa-Schuch, D., Silva-Garcia, M., Pilla, E., Engelke, M., Kilisch, M., Lenz, C., Moller, U., Nakamura, F., Urlaub, H. and Geiss-Friedlander, R. (2016) ‘DPP9 is a novel component of the N-end rule pathway targeting the tyrosine kinase Syk’, Elife, 5.

Karolchik, D., Hinrichs, A. S., Furey, T. S., Roskin, K. M., Sugnet, C. W., Haussler, D. and Kent, W. J. (2004) ‘The UCSC Table Browser data retrieval tool’, Nucleic Acids Res, 32(Database issue), pp. D493–6.

Katic, I., Xu, L. and Ciosk, R. (2015) ‘CRISPR/Cas9 Genome Editing in Caenorhabditis elegans: Evaluation of Templates for Homology-Mediated Repair and Knock-Ins by Homology-Independent DNA Repair’, G3 (Bethesda), 5(8), pp. 1649–56.

Kawasaki, I., Shim, Y. H., Kirchner, J., Kaminker, J., Wood, W. B. and Strome, S. (1998) ‘PGL-1, a predicted RNA-binding component of germ granules, is essential for fertility in C. elegans’, Cell, 94(5), pp. 635–45.

Keane, F. M., Nadvi, N. A., Yao, T. W. and Gorrell, M. D. (2011) ‘Neuropeptide Y, B-type natriuretic peptide, substance P and peptide YY are novel substrates of fibroblast activation protein-alpha’, FEBS J, 278(8), pp. 1316–32.

Kent, W. J., Zweig, A. S., Barber, G., Hinrichs, A. S. and Karolchik, D. (2010) ‘BigWig and BigBed: enabling browsing of large distributed datasets’, Bioinformatics, 26(17), pp. 2204–7.

Ketting, R. F., Haverkamp, T. H., van Luenen, H. G. and Plasterk, R. H. (1999) ‘Mut-7 of C. elegans, required for transposon silencing and RNA interference, is a homolog of Werner syndrome helicase and RNaseD’, Cell, 99(2), pp. 133–41.

Kluyver, T., Ragan-Kelley, B., Pérez, F., Granger, B. E., Bussonnier, M., Frederic, J., Kelley, K., Hamrick, J. B., Grout, J., Corlay, S. and Others ‘Jupyter Notebooks-a publishing format for reproducible computational workflows’, ELPUB, 2016, 87–90.

Koster, J. and Rahmann, S. (2018) ‘Snakemake-a scalable bioinformatics workflow engine’, Bioinformatics, 34(20), pp. 3600.

Kwak, P. B. and Tomari, Y. (2012) ‘The N domain of Argonaute drives duplex unwinding during RISC assembly’, Nat Struct Mol Biol, 19(2), pp. 145–51.

L’Hernault, S. W. (2009) ‘The genetics and cell biology of spermatogenesis in the nematode C. elegans’, Mol Cell Endocrinol, 306(1-2), pp. 59–65.

Langmead, B., Trapnell, C., Pop, M. and Salzberg, S. L. (2009) ‘Ultrafast and memory-efficient alignment of short DNA sequences to the human genome’, Genome Biol, 10(3), pp. R25.

Lewis, A., Berkyurek, A. C., Greiner, A., Sawh, A. N., Vashisht, A., Merrett, S., Flamand, M. N., Wohlschlegel, J., Sarov, M., Miska, E. A. and Duchaine, T. F. (2020) ‘A Family of Argonaute-Interacting Proteins Gates Nuclear RNAi’, Mol Cell, 78(5), pp. 862–875 e8.

Li, H., Handsaker, B., Wysoker, A., Fennell, T., Ruan, J., Homer, N., Marth, G., Abecasis, G., Durbin, R. and Genome Project Data Processing, S. (2009) ‘The Sequence Alignment/Map format and SAMtools’, Bioinformatics, 25(16), pp. 2078–9.

Liao, Y., Smyth, G. K. and Shi, W. (2014) ‘featureCounts: an efficient general purpose program for assigning sequence reads to genomic features’, Bioinformatics, 30(7), pp. 923–30.

MacLean, B., Tomazela, D. M., Shulman, N., Chambers, M., Finney, G. L., Frewen, B., Kern, R., Tabb, D. L., Liebler, D. C. and MacCoss, M. J. (2010) ‘Skyline: an open source document editor for creating and analyzing targeted proteomics experiments’, Bioinformatics, 26(7), pp. 966–8.

Madeira, F., Park, Y. M., Lee, J., Buso, N., Gur, T., Madhusoodanan, N., Basutkar, P., Tivey, A. R. N., Potter, S. C., Finn, R. D. and Lopez, R. (2019) ‘The EMBL-EBI search and sequence analysis tools APIs in 2019’, Nucleic Acids Res, 47(W1), pp. W636–W641.

Martin, M. (2011) ‘Cutadapt removes adapter sequences from high-throughput sequencing reads’, EMBnet.journal, 17(1), pp. 10–12.

McKinney, W. (2011) ‘pandas: a foundational Python library for data analysis and statistics’, Python for High Performance and Scientific Computing, pp. 1–9.

Mentlein, R. (1999) ‘Dipeptidyl-peptidase IV (CD26)--role in the inactivation of regulatory peptides’, Regul Pept, 85(1), pp. 9–24.

Mentlein, R., Gallwitz, B. and Schmidt, W. E. (1993) ‘Dipeptidyl-peptidase IV hydrolyses gastric inhibitory polypeptide, glucagon-like peptide-1(7-36)amide, peptide histidine methionine and is responsible for their degradation in human serum’, Eur J Biochem, 214(3), pp. 829–35.

Miki, T. S., Carl, S. H., Stadler, M. B. and Grosshans, H. (2016) ‘XRN2 Autoregulation and Control of Polycistronic Gene Expresssion in Caenorhabditis elegans’, PLoS Genet, 12(9), pp. e1006313.

Ostapcuk, V., Mohn, F., Carl, S. H., Basters, A., Hess, D., Iesmantavicius, V., Lampersberger, L., Flemr, M., Pandey, A., Thoma, N. H., Betschinger, J. and Buhler, M. (2018) ‘Activity-dependent neuroprotective protein recruits HP1 and CHD4 to control lineage-specifying genes’, Nature, 557(7707), pp. 739–743.

Padeken, J., Zeller, P., Towbin, B., Katic, I., Kalck, V., Methot, S. P. and Gasser, S. M. (2019) ‘Synergistic lethality between BRCA1 and H3K9me2 loss reflects satellite derepression’, Genes Dev, 33(7-8), pp. 436–451.

Phillips, C. M., Brown, K. C., Montgomery, B. E., Ruvkun, G. and Montgomery, T. A. (2015) ‘piRNAs and piRNA-Dependent siRNAs Protect Conserved and Essential C. elegans Genes from Misrouting into the RNAi Pathway’, Dev Cell, 34(4), pp. 457–65.

Phillips, C. M., Montgomery, T. A., Breen, P. C. and Ruvkun, G. (2012) ‘MUT-16 promotes formation of perinuclear mutator foci required for RNA silencing in the C. elegans germline’, Genes Dev, 26(13), pp. 1433–44.

Quinlan, A. R. and Hall, I. M. (2010) ‘BEDTools: a flexible suite of utilities for comparing genomic features’, Bioinformatics, 26(6), pp. 841–2.

Robinson, M. D., McCarthy, D. J. and Smyth, G. K. (2010) ‘edgeR: a Bioconductor package for differential expression analysis of digital gene expression data’, Bioinformatics, 26(1), pp. 139–40.

Sakaguchi, A., Sarkies, P., Simon, M., Doebley, A. L., Goldstein, L. D., Hedges, A., Ikegami, K., Alvares, S. M., Yang, L., LaRocque, J. R., Hall, J., Miska, E. A. and Ahmed, S. (2014) ‘Caenorhabditis elegans RSD-2 and RSD-6 promote germ cell immortality by maintaining small interfering RNA populations’, Proc Natl Acad Sci U S A, 111(41), pp. E4323–31.

Seth, M., Shirayama, M., Gu, W., Ishidate, T., Conte, D., Jr. and Mello, C. C. (2013) ‘The C. elegans CSR-1 argonaute pathway counteracts epigenetic silencing to promote germline gene expression’, Dev Cell, 27(6), pp. 656–63.

Seydoux, G. (2018) ‘The P Granules of C. elegans: A Genetic Model for the Study of RNA-Protein Condensates’, J Mol Biol, 430(23), pp. 4702–4710.

Shen, E. Z., Chen, H., Ozturk, A. R., Tu, S., Shirayama, M., Tang, W., Ding, Y. H., Dai, S. Y., Weng, Z. and Mello, C. C. (2018) ‘Identification of piRNA Binding Sites Reveals the Argonaute Regulatory Landscape of the C. elegans Germline’, Cell, 172(5), pp. 937–951 e18.

Smibert, P., Yang, J. S., Azzam, G., Liu, J. L. and Lai, E. C. (2013) ‘Homeostatic control of Argonaute stability by microRNA availability’, Nat Struct Mol Biol, 20(7), pp. 789–95.

Spracklin, G., Fields, B., Wan, G., Becker, D., Wallig, A., Shukla, A. and Kennedy, S. (2017) ‘The RNAi Inheritance Machinery of Caenorhabditis elegans’, Genetics, 206(3), pp. 1403–1416.

Tabara, H., Sarkissian, M., Kelly, W. G., Fleenor, J., Grishok, A., Timmons, L., Fire, A. and Mello, C. C. (1999) ‘The rde-1 gene, RNA interference, and transposon silencing in C. elegans’, Cell, 99(2), pp. 123–32.

Tinevez, J. Y., Perry, N., Schindelin, J., Hoopes, G. M., Reynolds, G. D., Laplantine, E., Bednarek, S. Y., Shorte, S. L. and Eliceiri, K. W. (2017) ‘TrackMate: An open and extensible platform for single-particle tracking’, Methods, 115, pp. 80–90.

Tyanova, S., Temu, T., Sinitcyn, P., Carlson, A., Hein, M. Y., Geiger, T., Mann, M. and Cox, J. (2016) ‘The Perseus computational platform for comprehensive analysis of (prote)omics data’, Nat Methods, 13(9), pp. 731–40.

Van Der Walt, S., Colbert, S. C. and others (2011) ‘The NumPy array: a structure for efficient numerical computation’, Comput. Sci. Eng.

Vasale, J. J., Gu, W., Thivierge, C., Batista, P. J., Claycomb, J. M., Youngman, E. M., Duchaine, T. F., Mello, C. C. and Conte, D., Jr. (2010) ‘Sequential rounds of RNA-dependent RNA transcription drive endogenous small-RNA biogenesis in the ERGO-1/Argonaute pathway’, Proc Natl Acad Sci U S A, 107(8), pp. 3582–7.

Vastenhouw, N. L., Fischer, S. E., Robert, V. J., Thijssen, K. L., Fraser, A. G., Kamath, R. S., Ahringer, J. and Plasterk, R. H. (2003) ‘A genome-wide screen identifies 27 genes involved in transposon silencing in C. elegans’, Curr Biol, 13(15), pp. 1311–6.

Villhauer, E. B., Brinkman, J. A., Naderi, G. B., Burkey, B. F., Dunning, B. E., Prasad, K., Mangold, B. L., Russell, M. E. and Hughes, T. E. (2003) ‘1-[[(3-hydroxy-1-adamanty-1-amino]acetyl]-2-cyano-(S)-pyrrolidine: a potent, selective, and orally bioavailable dipeptidyl peptidase IV inhibitor with antihyperglycemic properties’, J Med Chem, 46(13), pp. 2774–89.

Waskom, M., Botvinnik, O., Hobson, P., Warmenhoven, J., Cole, J. B., Halchenko, Y., Vanderplas, J., Hoyer, S., Villalba, S., Quintero, E. and Others (2014) ‘Seaborn: statistical data visualization’, URL: https://seaborn.pydata.org/(visited on 2017-05-15).

Wedeles, C. J., Wu, M. Z. and Claycomb, J. M. (2013) ‘Protection of germline gene expression by the C. elegans Argonaute CSR-1’, Dev Cell, 27(6), pp. 664–71.

Wedeles, C. J., Wu, M. Z. and Claycomb, J. M. (2014) ‘Silent no more: Endogenous small RNA pathways promote gene expression’, Worm, 3, pp. e28641.

Wilson, C. H., Zhang, H. E., Gorrell, M. D. and Abbott, C. A. (2016) ‘Dipeptidyl peptidase 9 substrates and their discovery: current progress and the application of mass spectrometry-based approaches’, Biol Chem, 397(9), pp. 837–56.

Wingfield, P. T. (2017) ‘N-Terminal Methionine Processing’, Curr Protoc Protein Sci, 88, pp. 6 14 1–6 14 3.

Wu, J. J., Tang, H. K., Yeh, T. K., Chen, C. M., Shy, H. S., Chu, Y. R., Chien, C. H., Tsai, T. Y., Huang, Y. C., Huang, Y. L., Huang, C. H., Tseng, H. Y., Jiaang, W. T., Chao, Y. S. and Chen, X. (2009) ‘Biochemistry, pharmacokinetics, and toxicology of a potent and selective DPP8/9 inhibitor’, Biochem Pharmacol, 78(2), pp. 203–10.

Yigit, E., Batista, P. J., Bei, Y., Pang, K. M., Chen, C. C., Tolia, N. H., Joshua-Tor, L., Mitani, S., Simard, M. J. and Mello, C. C. (2006) ‘Analysis of the C. elegans Argonaute family reveals that distinct Argonautes act sequentially during RNAi’, Cell, 127(4), pp. 747–57.

Youngman, E. M. and Claycomb, J. M. (2014) ‘From early lessons to new frontiers: the worm as a treasure trove of small RNA biology’, Front Genet, 5, pp. 416.

Zeller, P., Padeken, J., van Schendel, R., Kalck, V., Tijsterman, M. and Gasser, S. M. (2016) ‘Histone H3K9 methylation is dispensable for Caenorhabditis elegans development but suppresses RNA:DNA hybrid-associated repeat instability’, Nat Genet, 48(11), pp. 1385–1395.

Zhang, H., Chen, Y., Keane, F. M. and Gorrell, M. D. (2013) ‘Advances in understanding the expression and function of dipeptidyl peptidase 8 and 9’, Mol Cancer Res, 11(12), pp. 1487–96.

